# Severe Central Nervous System Demyelination in Sanfilippo Disease

**DOI:** 10.1101/2023.04.12.536631

**Authors:** Mahsa Taherzadeh, Erjun Zhang, Irene Londono, Sheng-Kwei Song, Sophie Wang, Jonathan D. Cooper, Timothy E. Kennedy, Carlos R. Morales, Zesheng Chen, Gregory A. Lodygensky, Alexey V. Pshezhetsky

## Abstract

Neurodegeneration and chronic progressive neuroinflammation are well-documented in neurological lysosomal storage diseases, including Sanfilippo disease or mucopolysaccharidosis III (MPS III). Since chronic neuroinflammation has been linked to white matter tract pathology and defects in axonal transmission, we analysed axonal myelination and white matter density in the mouse model of MPS IIIC and human post-mortem brain samples from MPS IIIA, C, and D patients. Analyses of corpus callosum (CC) and spinal cord tissues by immunohistochemistry revealed substantially reduced levels of myelin-associated proteins including Myelin Basic Protein, Myelin Associated Glycoprotein, and Myelin Oligodendrocyte Glycoprotein. Furthermore, ultrastructural analyses revealed disruption of myelin sheath organization and reduced myelin thickness in the brains of MPS IIIC mice and human MPS IIIC patients compared to healthy controls. Oligodendrocytes (OLs) in the CC of MPS IIIC mice were scarce, while examination of the remaining cells revealed numerous enlarged lysosomes containing heparan sulfate, GM3 ganglioside or “zebra bodies” consistent with accumulation of lipids and myelin fragments. In addition, OLs contained swollen mitochondria with largely dissolved cristae, resembling those previously identified in the dysfunctional neurons of MPS IIIC mice. When brains of 7-month-old MPS IIIC mice were analysed by ex-vivo Diffusion Basis Spectrum Imaging to assess microarchitectural changes in the corpus callosum, we found compelling signs of demyelination (26% increase in radial diffusivity) and tissue loss (76% increase in hindered diffusivity). Our findings demonstrate an import role for white matter injury in the pathophysiology of MPS III. Moreover, this study reveals specific parameters and brain regions for MRI analysis, a crucial non-invasive method to evaluate disease progression and therapeutic response in neurological lysosomal storage diseases.

## Introduction

About two-thirds of patients diagnosed with Lysosomal Storage Diseases (LSDs), a group of inherited metabolic disorders affecting lysosomal catabolism, show neurological symptoms and/or have pathological changes in the central nervous system (CNS) (reviewed in [1, 2]). Two of such disorders, Krabbe disease, caused by deficiency of β-galactocerebrosidase (GALC), and myeloid leukodystrophy (MLD), caused by deficiency of arylsulfatase A (ARSA), belong to the class of leukodystrophies, manifesting with fundamental abnormalities in the CNS, including white matter pathology and progressive degeneration of myelin sheaths (reviewed in [3]). Progressive myelin defects in these disorders are believed to be caused by two specific sphingolipids accumulating in the brains of affected patients, galactosylceramide also known as psychosine in the case of Krabbe disease and sulfatide in the case of MLD (reviewed in [3]). Both galactosylceramide and sulfatide are important components of myelin sheaths and are generated by myelin-producing cells, oligodendrocytes (OLs) in CNS, and Schwann cells in the peripheral nervous system. However, when the levels of galactosylceramide and sulfatide are drastically increased, as a result of genetic deficiencies of GALC and ARSA, respectively, they become highly toxic for OLs and Schwann cells. Because of progressive myelin loss, both Krabbe and MLD patients, especially those having an infantile-onset form of the disease caused by complete or almost complete deficiencies of the enzymes involved, manifest with a severe neurological impairment and deterioration leading ultimately to death before the age of 5 years [4–6]. In both Krabbe disease and MLD, MRI brain imaging was especially instrumental for the identification and characterization of white matter defects. In particular, extensive white matter defects were found in centrum semiovale, corpus callosum, and middle cerebellar peduncles of Krabbe patients by diffuse tension imaging (DTI) [7]. In infantile MLD patients, hyperintensity of T2-weighted MRI signals was observed in corpus callosum and in parieto-occipital regions suggesting defects in periventricular and central white matter [8]. Signs of demyelination (or delayed myelination) have been also detected by MRI in patients with multiple sulfatase deficiency, most likely due to the secondary deficiency of ARSA and the storage of sulfatide [9].

Since glycosphingolipids are critical components of myelin sheaths [10, 11], demyelination and white matter pathology have been also reported in patients affected with lysosomal glycosphingolipidoses, including Fabry disease, neurological forms of Gaucher disease (reviewed in [3]) and, most recently, infantile forms of Niemann-Pick disease type C (NPC) [12]. In the latter case, demyelination was proposed to results from the secondary storage of GM3 ganglioside in the lysosomes of OLs leading to their dysfunction [13]. White matter abnormalities have also been reported in patients diagnosed with other LSDs involving ganglioside accumulation, GM1 gangliosidosis [14] and GM2 gangliosidosis/Tay-Sachs disease [15].

Mucopolysaccharidosis type III (MPS III) or Sanfilippo disease, remains the most prevalent untreatable neurological lysosomal disorder (reviewed in [16]). MPS III is a spectrum of four conditions (MPS IIIA-D), caused by defects in the genes encoding the enzymes involved in lysosomal degradation of a glucosamine, heparan sulfate (HS). In MPS III, HS accumulates in brain tissue and causes neuronal dysfunction and death leading to neuropsychiatric problems, developmental delays, childhood dementia, blindness and death during the second decade of life (reviewed in [16]). MPS IIIA was found to be caused by defects in N-sulfoglucosamine sulfohydrolase (SGSH) [17], MPS IIIB, by defects in N-acetyl-α-D-glucosaminidase (NAGLU) [18], MPS IIIC, by defects in acetyl-CoA:alpha-glucosaminide N-acetyltransferase (HGSNAT) [19], and MPS IIID, by defects in N-acetylglucosamine-6-sulfate sulfatase (GNS) [20].

Previous studies involving MPS III patients and animal models revealed that disrupted catabolism of HS in the brain causes multifaceted pathological response, from neuroinflammation and oxidative stress to impairment of autophagy, neuronal accumulation of misfolded proteins, GM2 or GM3 gangliosides, and synaptic defects. Together, these processes contribute to neuronal dysfunction and neurodegeneration which lead to cognitive and motor decline in patients (reviewed in [16]). Importantly, in most Sanfilippo patients the rate of progressive functional impairment correlates with that of cortical and cerebellar atrophy, ventricular volume increase and other brain abnormalities, detected by computed tomography-scan (CT-scan) or magnetic resonance imaging (MRI), suggesting that these symptoms are associated with neuronal loss and pathological grey matter lesions (reviewed in [21, 22]). Much less is known, however, about the white matter defects and, in particular, axonal pathology and demyelination in the MPS III patients. While some case reports have indicated the presence of diffuse high-intensity signal in the white matter of MPS IIIA and IIIB patients [23, 24], other studies reported an absence of white-matter lesions in most MPS III cases [25].

Here, for the first time, we report that progressive severe demyelination is a hallmark of CNS pathology in both human MPS IIIC patients and the mouse *Hgsnat^P304L^* model [26] of the disease. We also demonstrate that, in the brains of MPS IIIC mice, the disruption of myelination results from reduced OL numbers and substantial pathological changes in OL morphology.

## Materials and Methods

### Study approval

All animal experiments were approved by the Animal Care and Use of CHU Ste-Justine Research Ethics Committee (approval numbers 2020-2658 and 2022-3452) and performed in accordance with the Canadian Council on Animal Care guidelines. Ethical approval for the research involving human tissues was granted by Research Ethics Board of CHU Ste-Justine.

### Animals

The knock-in mouse model of MPS IIIC, *Hgsnat^P304L^*, expressing the HGSNAT enzyme with human missense mutation P304L, on a C57BL/6J genetic background, has been previously described [26]. The Thy1-EGFP transgene mice expressing enhanced green fluorescent protein (EGFP) under the control of a modified regulatory region of the mouse thy1.2 gene promoter (containing sequences required for neuronal expression but lacking sequences required for expression in non-neural cells) were obtained from The Jackson Laboratory (JAX stock #007788) [27].

Mouse studies were performed in accordance with the Canadian Council on Animal Care (CCAC) in the CCAC-accredited animal facility of the CHU Ste-Justine. All mice were housed in an enriched environment in poly-carbonate cages under 12:12 h light:dark cycles in a temperature-and humidity-controlled room. Mice had access to a normal rodent chow and water *ad libitum*. The animals were bred as homozygous couples for both WT and *Hgsnat^P304L^* strains. Experiments were conducted using both male and female mice and the data analyzed to identify potential differences between sexes. Since no differences between sexes were observed in the experiments, the data for males and females were combined.

### Analysis of human brain tissues

Post-mortem brain tissues from two patients with MPS IIIC, fixed in 10% buffered formalin, were collected for neuropathological examination. The first brain was provided by the Anatomic Gift Registry while the second brain was donated for research purposes by the parents of the patient. Brain tissues of age-matched patients without CNS pathology were provided, together with clinical descriptions, by the Neuromax biobank of CHU Sainte-Justine.

The brains underwent gross examination by a neuropathologist (ZC), and representative regions were sampled and processed for paraffin embedding. Four-μm sections were cut from the formalin fixed paraffin embedded (FFPE) blocks, stained with Haematoxylin-Eosin (H&E), Saffron (HES), periodic acid-Schiff stain (PAS) and Luxol Fast Blue (LFB), and examined under the light microscope.

In addition, frozen or paraformaldehyde (PFA) fixed cerebral cortices from MPS patients (1 case of MPSIIIA, 1 case of MPSIIIC and 1 case of MPSIIID) and age-matched controls with no pathological changes in the central nervous system were provided by NIH NeuroBioBank (project 1071, MPS Synapse). Upon arrival to the laboratory, the samples were embedded in Tissue-Tek® optimum cutting temperature (OCT, Sakura, USA) compound and stored at-80°C. 40 µm thick brain sections were cut and stored in cryopreservation buffer (0.05 M sodium phosphate buffer pH 7.4, 15% sucrose, 40% ethylene glycol) and stored at-20°C until immunohistochemical labelling.

### Immunohistochemistry

Mice were perfused with 4% PFA in PBS. Following perfusion, brains were isolated and post-fixed in 4% PFA in PBS overnight. Brains were further incubated in 30% sucrose for 48 h at 4°C, embedded in OCT, cut into sequential 40 µm-thick coronal cross-sections using a Cryostat Epredia CryoStar NX50, and then kept at-20°C in cryopreservation buffer (0.05 M sodium phosphate buffer pH 7.4, 15% sucrose, 40% ethylene glycol). For immunofluorescence analysis, brain slices were permeabilized and blocked with 5% bovine serum albumin (BSA), and 0.3% Triton X-100 in PBS for 2 h at room temperature. Sections were then incubated with primary antibodies, diluted with 1% BSA and 0.1% Triton X-100 in PBS at 4°C overnight. The antibodies and their dilutions are shown in the Table 1.

**Table 1.**
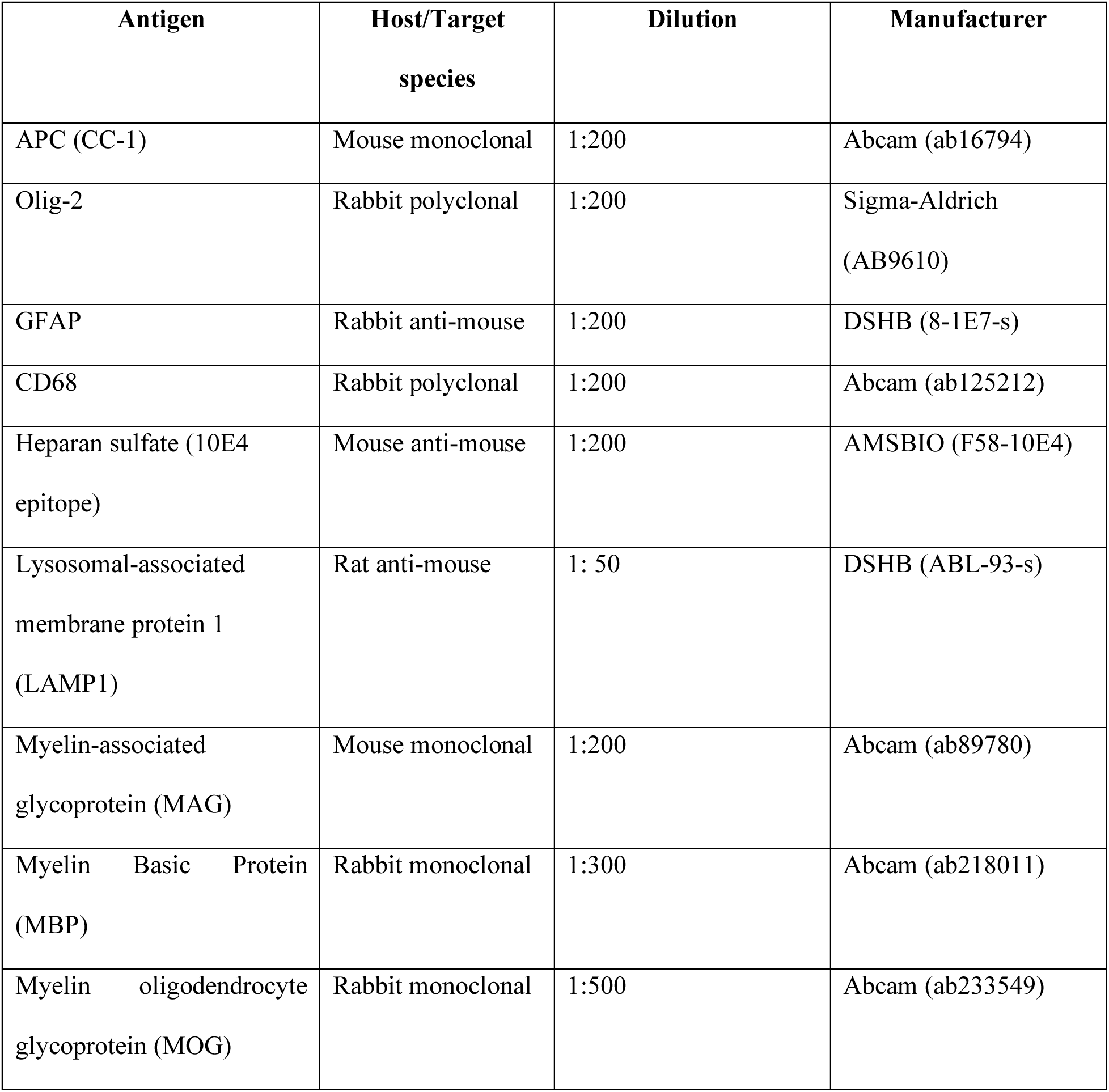
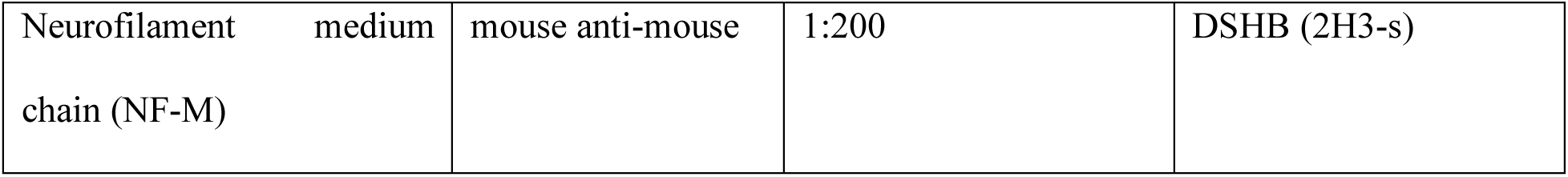
Antibodies and their dilutions used for immunochemistry

Mouse brain sections were washed with PBS and counterstained with Alexa Fluor® 488-conjugated goat anti-mouse IgG (A21202), Alexa Fluor® 555-conjugated goat anti-rabbit IgG (A21428), and Alexa Fluor® 633-conjugated goat anti-mouse IgG (A21094) (dilution 1:400, all from Thermo Fisher Scientific) for 2 h at room temperature. To quench autofluorescence, the mouse brain sections were dipped briefly in TrueBlack® Lipofuscin Autofluorescence Quencher (dilution 1:10, Biotium, 23007) for 1 min, and then washed with PBS. The slides were mounted with Prolong Gold Antifade mounting reagent with DAPI (Invitrogen, P36935) and captured using a Laser scanning confocal microscope (Leica TCS SP8: 20x, 40x, and 63x oil objective, N.A. 1.4). Z-stack projections were conducted with a step size of 0.3 µm to represent images.

For the analysis of spinal cord, adjacent one-in-forty-eight series of 40 µm coronal sections from each mouse were stained on slides using a modified immunofluorescence protocol [28, 29]. Sections were labeled with rat anti-CD68 (1:400, Bio-Rad MCA1957) or rat anti-MBP (1:500, Merck Millipore, MAB386) primary antibodies followed by AlexaFluor546 goat anti-rat (1:500, Invitrogen A11081) for CD68, or AlexaFluor488 goat anti-rat (1:500, Invitrogen A48262) for MBP. Slides were counterstained with TrueBlack Lipofuscin Autofluorescence Quencher (Biotium).

To quantify microgliosis (CD68-positive activated microglia) and myelination (MBP immunoreactivity), semiautomated thresholding image analysis was performed as described previously [28, 29]. This involved collecting slide-scanned images at 10x magnification (Zeiss Axio Scan Z1 Fluorescence Slide Scanner) from each animal. Contours of appropriate anatomical regions were then drawn and images were subsequently analyzed using *Image-Pro Premier* (Media Cybernetics) using an appropriate threshold that selected the foreground immunoreactivity above the background. All thresholding data (CD68 and MBP) were expressed as the percentage of area within each anatomically defined region of interest that contained immunoreactivity above the set threshold for that antigen (“% immunoreactivity”).

### Image processing and analysis

Unless indicated otherwise, images were processed and quantified using ImageJ 1.50i (National Institutes of Health, Bethesda, MD, USA). Quantification was performed in a double-blind fashion. Three-dimensional images were generated using Imaris (Oxford Instruments, version 9.6) software (Bitplane).

### Immunoblotting

Half-brain sections from 6-month-old mice were homogenized in a non-denaturing lysis buffer (50 mM Tris-HCl, pH 7.4, 150 mM NaCl, 1% NP-40, 0.25% sodium deoxycholate, 0.1% SDS, 2 mM EDTA, 1 mM PMSF), supplemented with protease and phosphate inhibitor cocktails (Sigma-Aldrich, cat# 4693132001 and 4906837001). The homogenates were cleared by centrifugation at 13,000 x g at 4°C for 25 min and the protein concentration in the collected supernatant was measured using a Pierce BCA Protein Assay Kit (Thermo Fisher Scientific). Protein extracts (20 µg of protein for each sample), were incubated in a boiling bath for 10 min and analyzed by SDS-PAGE on 4–20% precast polyacrylamide gradient gel (Bio-Rad, 4561096). Western blot analyses were performed according to standard protocols using antibodies against MBP (1:1000, Abcam,: cat# ab218011), MAG (1:1000, Abcam, cat# ab89780), and α-tubulin (1:2000, DSHB, cat# 12G10) as a control. The immunoblots were revealed by chemiluminescence with SuperSignalWest Pico PLUS (Thermo Fisher Scientific, Waltham, MA, USA). Detected bands were quantified using ImageJ 1.50i software (National Institutes of Health, Bethesda, MD, USA) and normalised for the intensity of the α-tubulin band.

### Transmission Electron Microscopy Analysis

To prepare sections for Transmission Electron Microscopy (TEM) analysis, 3 mice from each genotype were anesthetized with sodium pentobarbital (50 mg/kg BW) and perfused with phosphate-buffered saline (PBS, pH 7.4), followed by 2% paraformaldehyde / 2.5% glutaraldehyde in 0.1 M sodium cacodylate buffer, pH 7.4. The brains were incubated in the same fixative for 24 h at 4°C, washed with MilliQ H₂O, and post-fixed with 1% osmium tetroxide, 1.5% potassium ferrocyanide in H₂O for 2 h, at 4°C. The samples were further dehydrated in a graded bath of acetone and MilliQ H₂O with increasing concentrations of 30%, 50%, 70%, 80%, 90%, and 3 x 100% for 8 min. The samples were infiltrated with 100% Epon, embedded in a rubber embedding mold, and polymerized in the oven at 60°C for 48 hrs. Once the resin was polymerized, semi-thin (1.0 µm) sections of the corpus callosum were dissected and stained with 1% toluidine blue before being mounted on glass slides and examined with a Leica DMS light microscope to select regions of interest. The sections were cut into ultrathin (70–80 nm) sections, placed on 200 Mesh copper grids, stained with lead citrate, and examined at 80 kV on an FEI Tecnai G2 Spirit (FEI Company, Hillsboro, OR) electron microscope equipped with a Morada CCD digital camera (Olympus, Tokyo, Japan). Micrographs were taken with 2900 x and 4800 x magnification.

To quantify the number of myelinated axons, a minimum of 10 images were analyzed for each animal. All myelinated axons within the counting frame were counted and included in the statistical analysis. Then, for each micrograph, 5 randomly selected axons were used to analyze axonal parameters including axonal diameter, myelin thickness and g-ratio, using ImageJ software.

### Preclinical high-field MRI

Two groups of 7-month-old mice (10 WT and 8 MPS IIIC) were imaged using a preclinical scanner at the Cerebral Imaging Centre of the Douglas Mental Health University Institute (Montreal, Canada). Animals were perfused with PBS followed by 4% PFA in PBS, under terminal anesthesia, and brains were carefully removed and immersed in 4% PFA in PBS for 5 h. Brains were then mounted in a syringe with Fomblin oil for *ex vivo* MR imaging, using a solenoid coil custom-built to fit the syringe.

Imaging was carried out on a 7T Bruker MRI scanner. 2D spin-echo sequences with pulsed gradients were used to acquire diffusion-weighted data. Imaging included 1 b0 and 25 b-values (0 < b ≤ 3000 s/mm^2^) with different directions for each b-value, a field of view 12 mm × 12 mm, resolution, 0.15 × 0.15 × 0.4 mm^3^, and TR/TE: 3300 ms/32 ms. DBSI and DIPY-DTI (non-linear algorithm) were used for reconstruction as described [30, 31]. On two consecutive slices, regions of interest (ROI) were manually selected on corpus callosum in color-coded fractional anisotropy (FA) maps extracted from DTI.

### Statistical analysis

Statistical analyses were performed using Prism GraphPad 9.3.0. software (GraphPad Software San Diego, CA). The normality for all data was checked using the D’Agostino & Pearson omnibus normality test. Significance of the difference was determined using a Student’s t-test, when comparing two groups, and one-way ANOVA test followed by Tukey’s multiple comparison test, when comparing more than two groups. Two-way ANOVA followed by Bonferroni or Tukey’s post hoc tests was used for two-factor analysis. The Mann-Whitney test was used on metrics extracted from DBSI when normality was not attainted. A P-value less than 0.05 was considered significant.

## Results

### Massive reduction of myelin-associated proteins in MPS IIIC mice

Chronic progressive neuroinflammation and microgliosis (increased brain levels of proinflammatory activated microglia) are well documented in lysosomal storage diseases including MPS IIIC. As chronic neuroinflammation and microglial activation can alter white matter tracts and interrupt communication between neurons [32], we hypothesized that pathological changes in axon tracts might include a loss of myelin with decreased white matter density and contribute to CNS pathology in MPS IIIC patients. To investigate this hypothesis, we examined the integrity of myelination in our recently described *Hgsnat^P304L^*mouse model expressing the mutant HGSNAT Pro304Leu variant[26]. These mice reveal progressive pathological alterations in the cortical and subcortical gray matter, including pronounced synaptic defects, astromicrogliosis and neurodegeneration[26].

To assess for the presence of myelination defects, we first examined the levels of myelin-associated proteins, Myelin Basic Protein (MBP), Myelin Associated Glycoprotein (MAG) and Myelin Oligodendrocyte Glycoprotein (MOG), in the somatosensory cortices (SSC) and corpus callosum (CC) by immunohistochemical analysis of brain sections of 6-month-old *Hgsnat^P304L^* mice compared with WT mice matched for age, sex, and genetic background. The immunoreactivity detected for each protein was significantly reduced in the CC of MPS IIIC mice at 6 months of age compared with WT counterparts (Figure 1A-C). In SSC, levels of MAG were significantly reduced, while nonsignificant trends toward a decrease were observed for MBP and MOG (Figure 1D-F). The axonal marker, Neurofilament medium chain protein (NF-M), also showed a non-significant trend toward a decrease in both areas. Together, these results were consistent with white matter injury in the brain of 6-month-old *Hgsnat^P304L^*mice.

**Figure 1.**
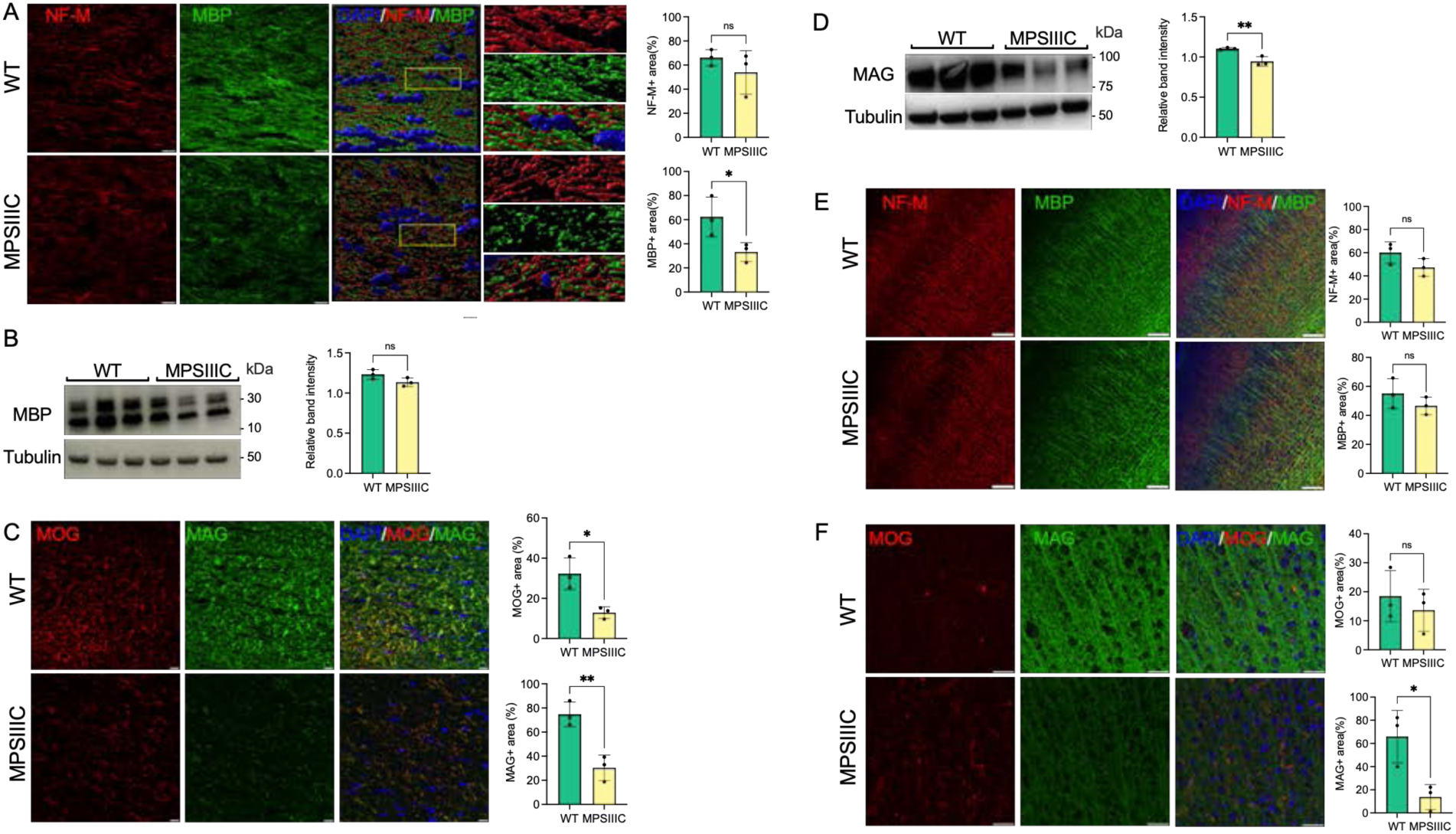
Myelin-associated proteins MBP, MAG, and MOG are reduced in the CC of MPSIIIC, compared with WT mice, suggesting myelin loss. (**A**) Panels show representative images of the CC of 6-month-old WT and MPSIIIC mice (left) and three-dimensional (3D) enlarged images of the areas marked by yellow boxes (right). MBP+ areas (green) on the surface of neurofilament medium chain (NF-M)+ axons (red) are reduced. The graphs show quantification of MBP+ and NF-M+ areas by ImageJ software. (**B**) Immunoblotting shows trend for reduction of MBP in the total brain homogenates of MPSIIIC mice. (**C**) IHC analysis reveals a reduction of MAG and MOG in CC of MPSIIIC mice compared with WT mice. Graphs show quantification of MAG+ and MOG+ stained area by ImageJ software. (**D**) Western blots of total protein extracts from brains of 6-month-old MPSIIIC mice confirm reduction of MAG. The graph shows intensities of MAG immunoreactive bands, quantified with ImageJ software and normalized by the intensity of tubulin immunoreactive bands. (**E**) No significant differences in MBP labelling in the cortex are detected by IHC between MPSIIIC and WT mice. (**F**) Level of MAG (green) labelling is reduced in the cortex of MPSIIIC compared with WT mice, while MOG (red) labelling shows only a non-significant trend for decrease. In all panels DAPI (blue) was used as a nuclear counterstain and scale bars equal 10 µm. All graphs show individual data, means and SD obtained for 3 mice per genotype. P-values were calculated using an unpaired t-test (*, p<0.05; **, p<0.01, ns, nonsignificant).

In the spinal cord of 6-month-old MPSIIIC mice, immunostaining for the microglial marker CD68 revealed numerous intensely stained microglia with enlarged cell soma in the grey and white matter, while age matched WT mice revealed only few lightly stained microglia with a small cell soma (Figure 2). Immunostaining for MBP revealed intense immunoreactivity within the spinal white matter of mice of both genotypes. Thresholding image analysis confirmed that significantly more CD68 immunoreactivity was present in both the dorsal and ventral grey matter of MPSIIIC mice. We detected a moderate yet significant reduction in the intensity of MBP immunoreactivity in the dorsal funiculus of MPSIIIC mice, but no significant difference in MBP immunoreactivity between genotypes in the ventral funiculus (Figure 2).

**Figure 2.**
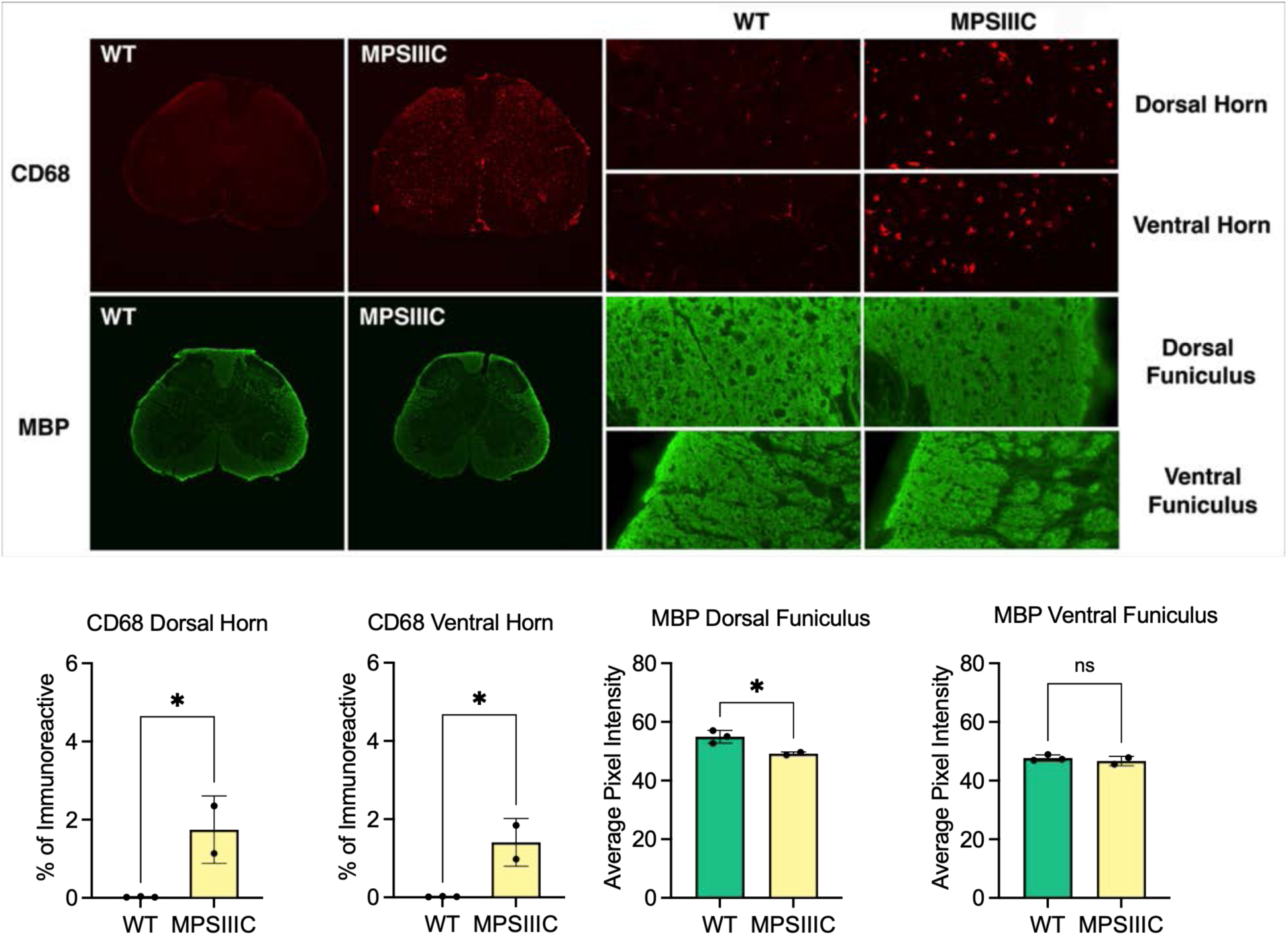
Microglial activation and white matter changes in the lumbar spinal cord of 6-month-old MPSIIIC mice. Representative images of immunostaining for the microglial marker CD68 (red) and myelin basic protein (MBP green) in the lumbar cord of 6-month-old *Hgsnat^P304L^* (MPSIIIC) mice and age-matched wild type (WT) control mice. Numerous CD68 immunoreactive microglia with enlarged cell soma are present in the gray and white matter of the dorsal and ventral horn of MPSIIIC mice but are virtually absent in WT mice at this age. Compared to age-matched control WT mice, MBP immunoreactivity is moderately reduced in the dorsal funiculus of 6-month-old MPSIIIC mice, but unchanged in the ventral funiculus. Quantitative thresholding image analysis confirmed these observations revealing significantly elevated CD68 immunoreactivity in the dorsal and ventral horn of 6-month-old HSGNAT KI vs. age-matched WT mice. Similarly there was significantly less MBP immunoreactivity in the dorsal funiculus of 6-month-old HSGNAT KI vs. age-matched WT mice, but no significant change in the ventral funiculus. All graphs show individual data, means and SD obtained for 2 mice per genotype. P-values were calculated using an unpaired t-test (*, p<0.05; **, p<0.01, ns, nonsignificant).

### Reduction of myelin thickness and axon degeneration in the CC of MPSIIIC mice

To determine if changes in the amounts of myelin associated proteins in the CC coincided with alterations in myelin structure, three MPSIIIC and three age/sex matched WT mice were examined at the ultrastructural level by TEM followed by quantification of axon diameters and myelin thickness. We detected a relative scarcity of myelinated axonal profiles in MPSIIIC mice, and the axons that were myelinated had an average g-ratio (axon diameter/myelinated fiber diameter) of 0.803 ± 0.134, that was significantly higher than the one for WT mice, 0.76 ± 0.15, indicating a reduced thicknesses of myelin sheaths (Figure 3). Scatter plots of g-ratios vs axonal diameter indicated hypomyelination of axons of all sizes in MPSIIIC mice. At the same time, no difference in the mean axonal diameters was detected between WT and mutant mice. Notably, in control animals, myelin thickness showed the expected positive correlation with axon diameter (R^2^ = 0.25), while, in MPSIIIC mice, myelin thickness did not significantly correlate with axon diameter (R^2^ = 0.08). TEM examination revealed defects in myelination along axons in CC (empty myelin sheath and splits in the compact myelin) with axonal swelling (spheroids) containing accumulated storage and/or transport vesicles. Similar swellings that coincided with microtubule defects were previously reported by us in hippocampal CA1 pyramidal neurons and in cultured neurons of another MPSIIIC model (*Hgsnat-Geo* mice) along with the evidence that the swellings disrupt synaptic vesicle precursor transport along axons[36].

**Figure 3.**
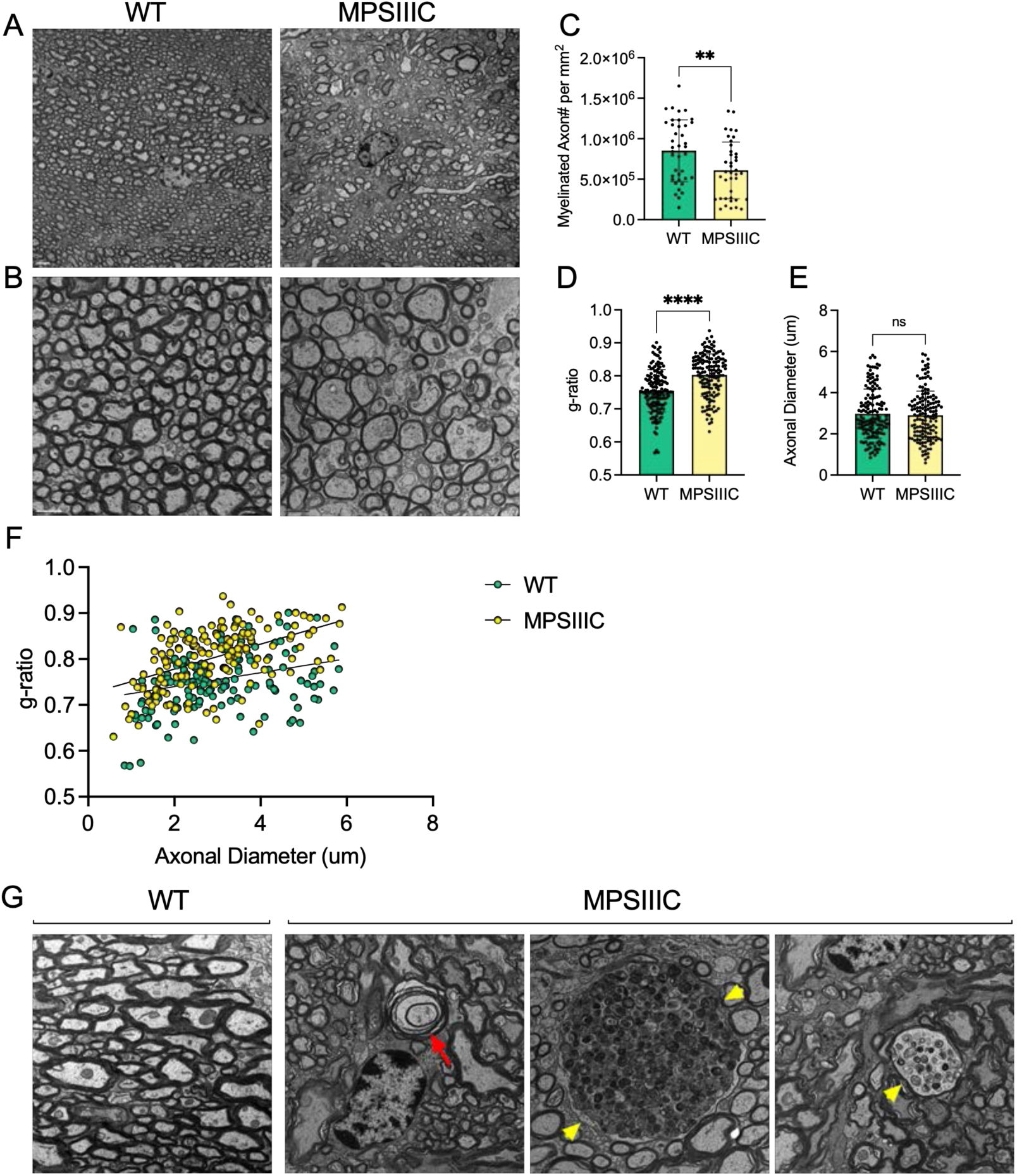
Transmission electron microscopy reveals reduced myelin thickness, decreased number of myelinated axons, structural defects in myelin sheaths and axonal swelling in the corpus callosum of MPSIIIC mouse. Panels (**A**) and (**B**) show representative TEM images of CC of 6-month-old MPSIIIC and WT mice taken at lower (2000X) and higher (4000X) magnifications, respectively. Scale bars equal 2 mm (**A**) and 1 mm (**B**). (**C**) The graph shows quantification of myelinated axon density (mean number of myelinated axons per square millimeter) in CC of WT and MPSIIIC mice. (**D**) G-ratio values for myelinated axons in CC of MPSIIIC mice are significantly higher than those for axons of WT mice. (**E**) Axonal diameters are similar in WT and MPSIIIC mice. (**F**) Scatter plot depicting g-ratio versus axonal diameter values. Graphs in panels (**D-E**) show individual values, means and SD and in the panel (**F**) individual values and linear regression plots. Sections from 3 mice per genotype (50 randomly selected axons per mouse) were analyzed. (**G**) Electron micrographs show an absence of pathological changes in the axons of WT mice. In contrast, degenerated axons with empty myelin sheath and split myelin (red arrow), as well as large axonal swellings containing accumulating vesicles (yellow arrowheads), are observed in MPSIIIC mice.

### Activated microglia in CC of MPSIIIC mice accumulate myelin debris

Myelin degradation involves its fragmentation with the resulting fragments being taken up by microglia[33, 34]. We, therefore, analyzed brain tissues of MPSIIIC mice using IHC to determine if microglia contained myelin fragments. MBP-positive puncta were readily detected in ILB4-positive (Figure 4A) and CD68-positive (Supplementary figure S1) microglia in the CC of MPS IIIC mice but not of WT mice. Moreover, MBP-positive puncta in microglia localized inside enlarged LAMP1-positive vacuoles, confirming lysosomal accumulation of MBP fragments in these cells. The IHC results were confirmed by TEM examination revealing that microglia in the CC of MPS IIIC mice were enlarged and contained vacuoles with so called “zebra bodies”, consistent with myelin accumulation (Figure 4B, boxed). We also detected electrolucent vacuoles most probably containing HS and other glycosaminoglycans (Figure 4B, arrows) [35]. In the WT mice, the microglia remained small and did not contain enlarged vacuoles (Figure 4B). Notably, CD68-positive activated microglia were detected in MPSIIIC brain as early as P25, when the levels of MBP in the CC are still intact (Supplementary figure S2), suggesting that demyelination was likely not caused by direct action of the immune cells.

**Figure 4.**
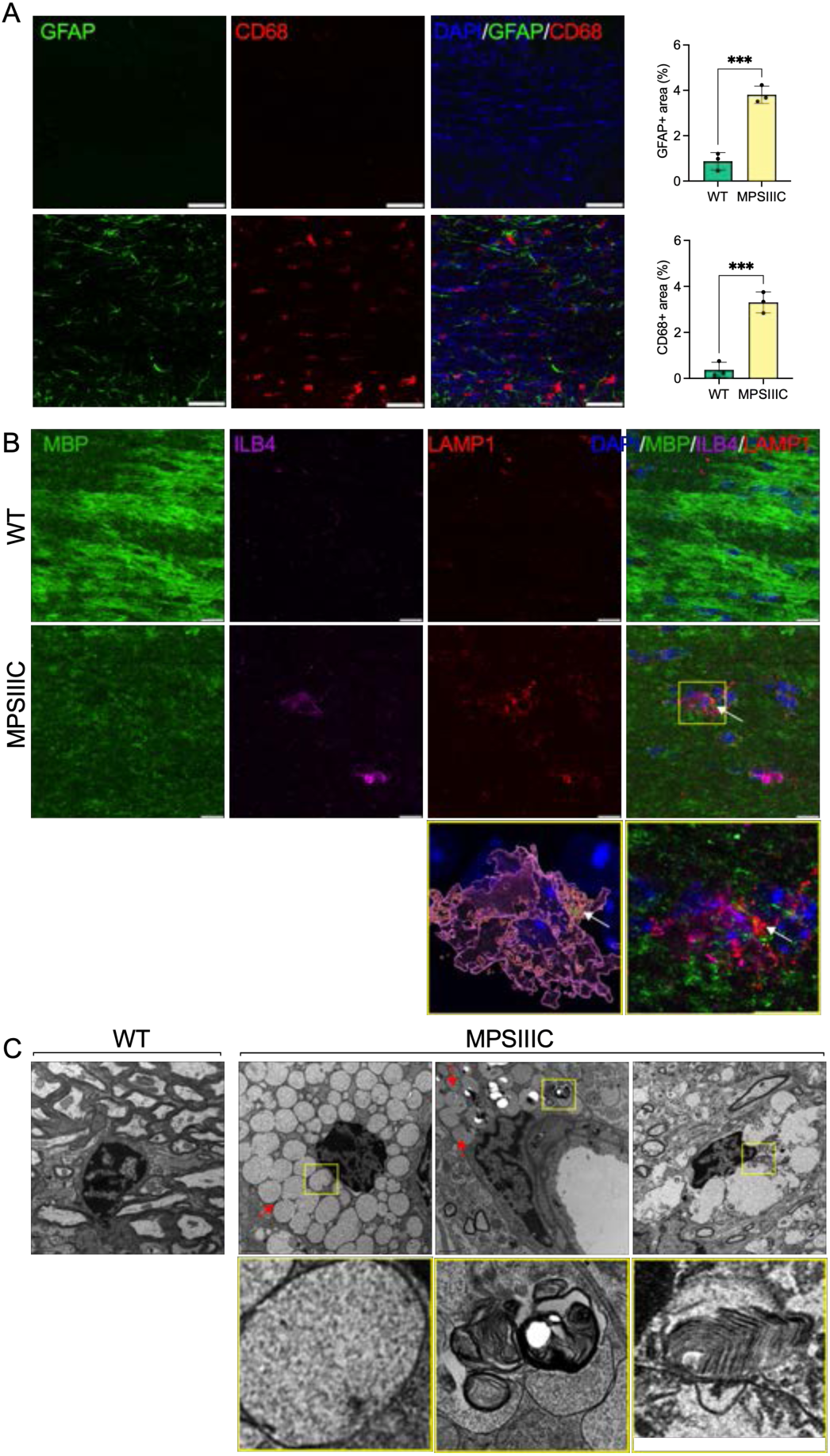
Microglia in CC of MPSIIIC but not of WT mice show lysosomal accumulation of myelin fragments and GAGs. **(A)** Panels show representative confocal microscopy images of CC tissue of 6-month-old WT and MPSIIIC mice labelled with antibodies against GFAP (green) and CD68 (red), markers for astrocytes and activated microglia, respectively. DAPI (blue) was used as a nuclear counterstain. Scale bar equals 50 µm. **(B)** Panels show representative confocal microscopy images of the CC of 6-month-old MPSIIIC and WT mice labelled with fluorescent isolectin b4 (ILB4, purple), and antibodies against MBP (green), and LAMP1 (red). DAPI (blue) was used as a nuclear counterstain. Scale bars equal 10 µm. The enlarged confocal image of the boxed area shows the colocalization of MBP+ puncta with LAMP1+ lysosomal marker inside ILB4+ activated microglia in the CC of a MPSIIIC mouse. 3D reconstruction shows that MBP+ puncta (arrows) are located inside the LAMP1+ lysosome of a microglia cell. **(C)** Both electron-lucent vacuoles (arrow) consistent with HS storage, and those containing “zebra bodies”, indicative of myelin debris (boxed), are detected in microglia in the CC of MPSIIIC mouse. Microglia in the CC of WT mouse are small and do not contain storage vacuoles. Scale bar equals 1 mm.

### Oligodendrocyte dysfunction in MPSIIIC mice

To understand the mechanism underlying demyelination, we analyzed the abundance and morphological phenotype of OLs in the CC of MPSIIIC and WT mice. To study oligodendrocyte maturation, we performed co-immunostaining of the brain tissues with antibodies against Olig2, a marker expressed both by oligodendrocytes (OLs) and oligodendrocyte precursor cells (OPCs), and antibodies against CC1, a marker of mature OLs. Our data (Figure 5A and Supplementary figure S3) showed that the majority of CC1-positive cells in the CC of WT mice were co-labelled for Olig2. On the other hand, in the brains of MPSIIIC mice, we found a significant reduction of CC1-positive, Oligo2-positive as well as Olig2/CC1 double-positive cells suggesting either the reduced production of OLs or their increased degeneration (Figure 5A and Supplementary figure S3).

**Figure 5.**
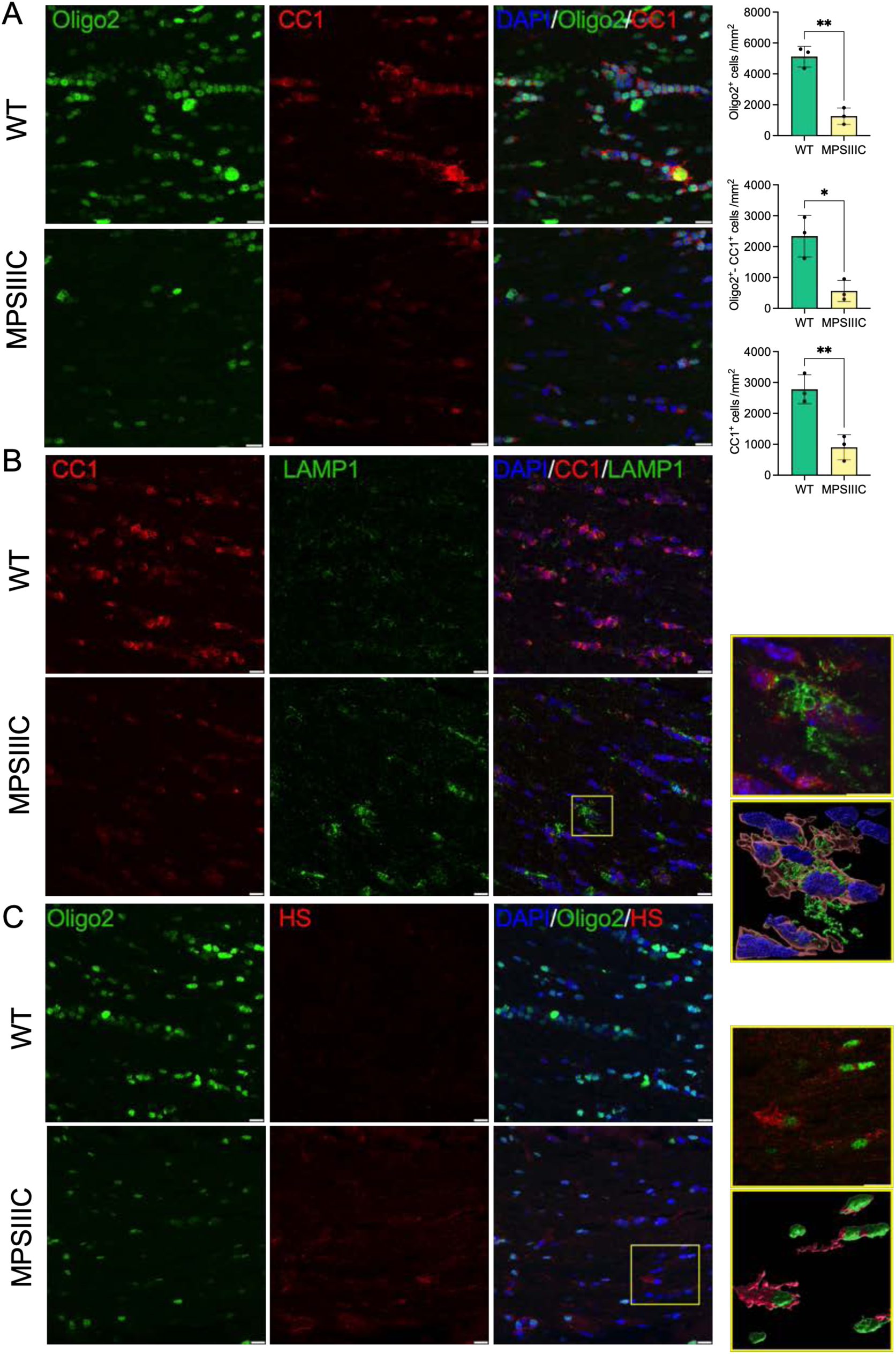
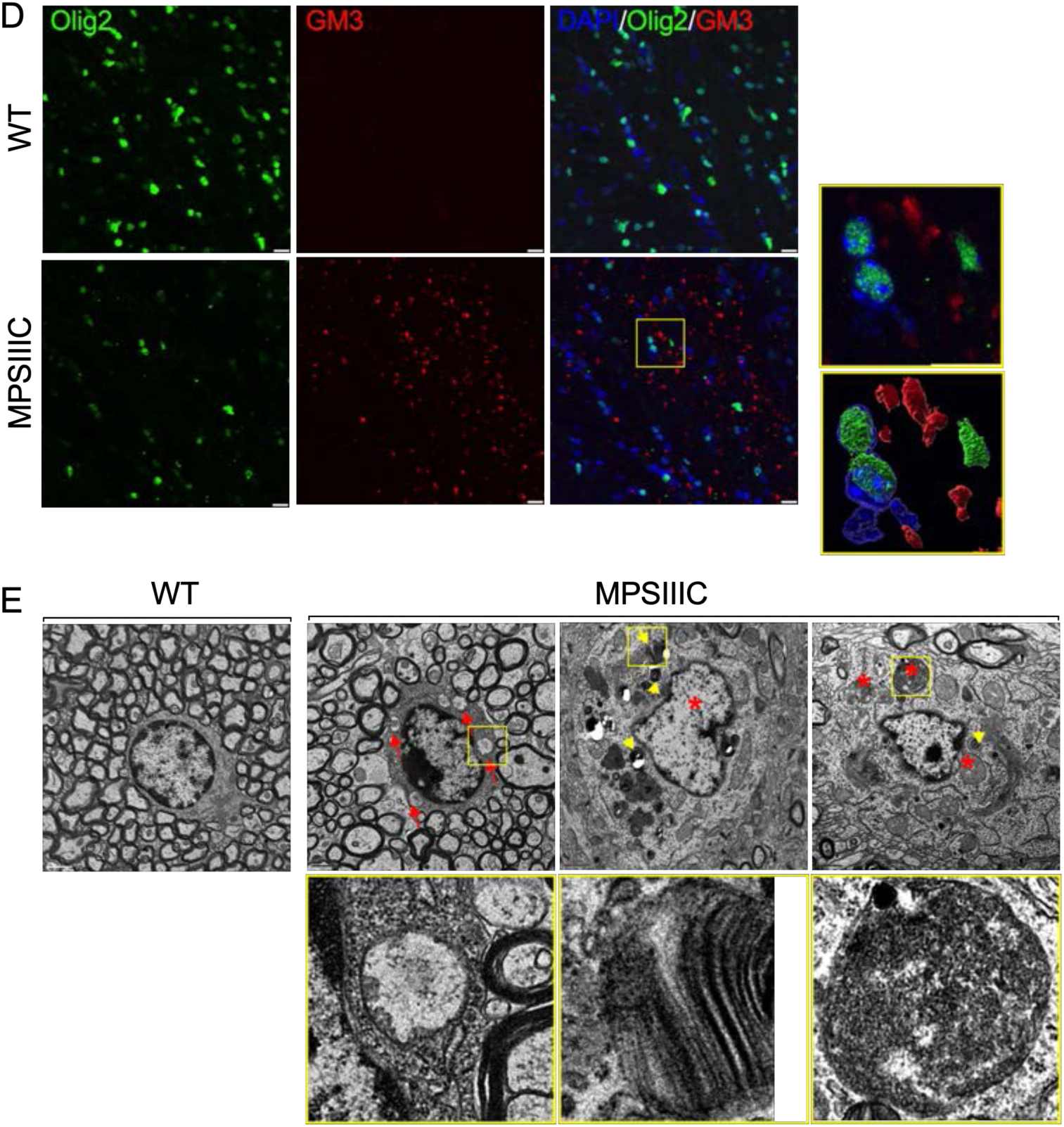
Oligodendrocytes in CC of MPSIIIC mice show reduced abundance, maturation and morphological abnormalities. **(A)** Representative confocal microscopy images of CC of 6-month-old WT and MPSIIIC mice immunolabelled for OL lineage marker Olig2 (green) and mature OL marker CC1 (red). Graphs show quantification of Oligo2+, CC1+ and Oligo2+/CC1+ cells (the number of cells per mm^2^) in WT and MPSIIIC mice. **(B)** Images of CC of 6-month-old WT and MPSIIIC mice immunolabelled for CC1 (red) and LAMP1 (green). Reconstructed 3D images of cells in the enlarged area boxed in the right panel show a significant increase in the size and abundance of LAMP1+ puncta consistent with presence of enlarged lysosomes in the OLs of MPSIIIC mice. **(C)** Representative images of CC of 6-month-old WT and MPSIIIC mice immunolabelled for Olig2 (green) and HS (red) show the accumulation of HS in the Olig2+ cells. **(D)** Images of CC of 6-month-old WT and MPSIIIC mice immunolabelled for Olig2 (green) and GM3 (red) show accumulation of GM3 ganglioside in the OLs of MPSIIIC mice. For all panels DAPI (blue) was used as a nuclear counterstain. Bars equal 10 µm. **(E)** TEM micrographs of OLs reveal numerous storage vacuoles, both electrolucent (red arrows) and those containing zebra bodies (yellow arrowheads), as well as swollen mitochondria with largely dissolved cristae (asterisks). High magnification images of boxed areas show a detailed view of storage deposits. Scale bars equal 1 µm. All graphs show individual data, means and SD obtained for 3 mice per genotype. P-values were calculated using an unpaired t-test.

The reduced number of OLs coincided with severe morphological changes observed in most of the remaining cells. Specifically, the number and size of LAMP1-positive vacuoles in OLs were increased, consistent with a lysosomal storage phenotype (Figure 5B). Our previous studies revealed that HGSNAT deficiency and impairment of HS catabolism resulted in intralysosomal accumulation of both primary (HS) and secondary (gangliosides) storage materials in neurons and microglia in the brains of MPSIIIC mice[35, 36]. To investigate if both compounds are also stored in OLs of MPSIIIC mice, brain tissues were studied by immunohistochemistry using the 10E4 monoclonal antibody, specific for a native HS epitope, and the anti-GM3 ganglioside monoclonal antibody. These experiments revealed increased storage of both HS and GM3 gangliosides in multiple Oligo2-positive cells (Figure 5C and Supplementary figure S4). TEM analysis further confirmed that OLs in the CC of MPSIIIC mice (identified by the shape and pattern of their nuclei and the presence of multiple microtubular structures in the cytoplasm [37]) contained multiple storage vacuoles. Some vacuoles exhibited an electrolucent content compatible with storage of HS [38] (marked with red arrowheads and boxed in Figure 5D), while others contained zebra bodies suggesting storage of myelin fragments or/and sphingolipids [39] (marked with yellow arrowheads and boxed in Figure 5D). In addition, OLs in the CC of MPSIIIC mice contained swollen mitochondria with largely dissolved cristae (marked with asterisks in Figure 5D), similar to those we have previously identified in the neurons of MPSIIIC mice [35].

### High-field magnetic resonance diffusion imaging analysis of MPSIIIC mice reveals microarchitectural changes in the corpus callosum compatible with demyelination

In the corpus callosum, DTI analysis identified clear signs of white matter injury with significant increases in Radial diffusivity (RD, 26%, p=0.003), an indicator of demyelination [40, 41], and Mean diffusivity (MD, 15%, p=0.02) a measure known to inversely correlate with white matter maturation (Figures 6). The DBSI analysis showed an increase in Hindered fraction (HF, 76%, p<0.01) and in Water fraction (WF, 134%, p<0.02), that revealed loss of tissue in the CC of MPSIIIC mouse brains compared to controls (Figure 6).

**Figure 6:**
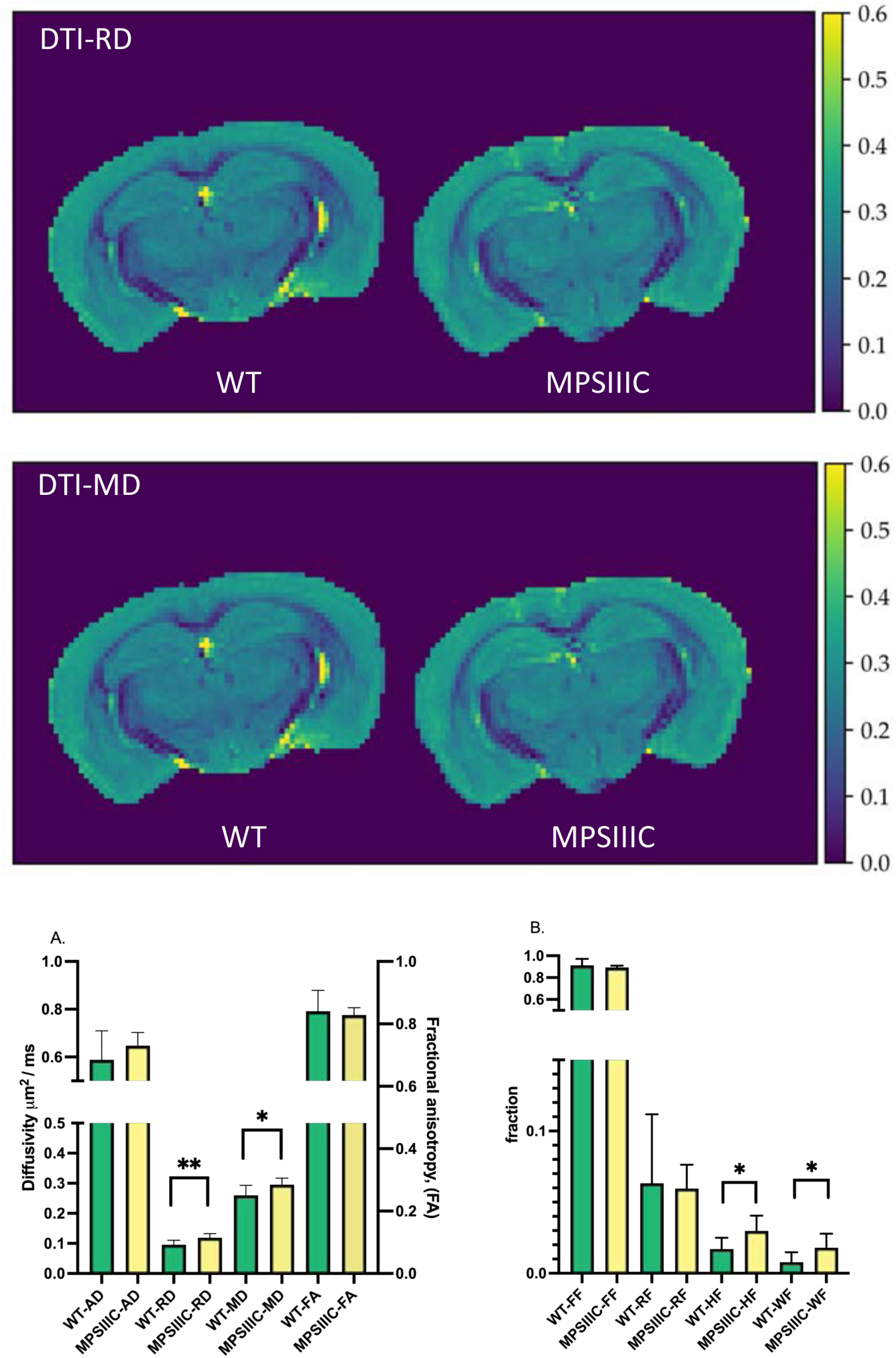
Diffusion maps from DTI and diffusion metrics from DTI and DBSI for WT and MPSIIIC mice. **(A)** Representative images of Radial diffusivity and Mean diffusivity maps for brains of WT and MPSIIIC mice demonstrating an increase for both parameters in MPSIIIC brain compared to WT mice. Arrows show the corpus callosum. Diffusivity scale (µm^2^/ms) is shown in the right sidebar. Abbreviations: RD, Radial diffusivity; MD, Mean diffusivity. **(B)** Bar plots of diffusion metrics from DTI (left) and DBSI (right) of WT and MPSIIIC mice. Abbreviations: RD, Radial diffusivity; MD, Mean diffusivity; HF, Hindered fraction, WF, Water fraction; Graphs show means ± SD for 8 WT (green bars) and 10 MPSIIIC (yellow bars) mice. **, p<0.01, * p<0.05.

### Myelination defects are pronounced in brain tissues of human MPSIII patients

We further analysed if axonal demyelination was also present in post-mortem tissues collected during autopsy of two MPSIIIC patients whose families provided informed consent for the use of the tissue in this research. The first patient was a 35-year-old Caucasian male with known diagnosis of MPSIIIC. He was wheelchair-dependent since the age of 15 and had the mental capacity of a 2-year-old. The second patient was an 18-year-old female with MPSIIIC diagnosed by biochemical assay of HGSNAT activity at the age of 7 years. The diagnosis was further confirmed by molecular analysis revealing that the patient was homozygous for the c.494-1G>A/p.[P164_S187delinsQSCYVTQAGVRWHHLGSLQALPPGFTPFSYLSLLSSWN, L165PfsX5] mutation affecting the conserved consensus sequence of the splice acceptor site in intron 4 [42].

On gross examination, the brain of the first patient showed severe cortical atrophy with fibrotic leptomeninges and enlarged lateral ventricles. The brain of the second patient had a marked decrease in brain weight of 974 g (normal: 1233 ± 115 g [43]) with severe cortical atrophy and enlarged lateral ventricles. However, it did not show any significant leptomeningeal fibrosis. On histological examination, both MPS brains showed neurons with abundant PAS-positive cytoplasmic inclusions (Figure 7A) at each of the three levels of the brain examined. These inclusions did not show LFB staining, except for the temporal lobe cortical neurons of the second patient (Figure 7B). HES sections of subcortical white matter in MPS brains showed slightly increased cellularity (Figure 7C, D) compared to the control brain. The first patient had a focal demyelinating lesion located at the anterior commissure with several hemosiderin-laden macrophages located in proximity (Figure 7E, F, G). LFB staining did not reveal any other regions with profound demyelination.

**Figure 7.**
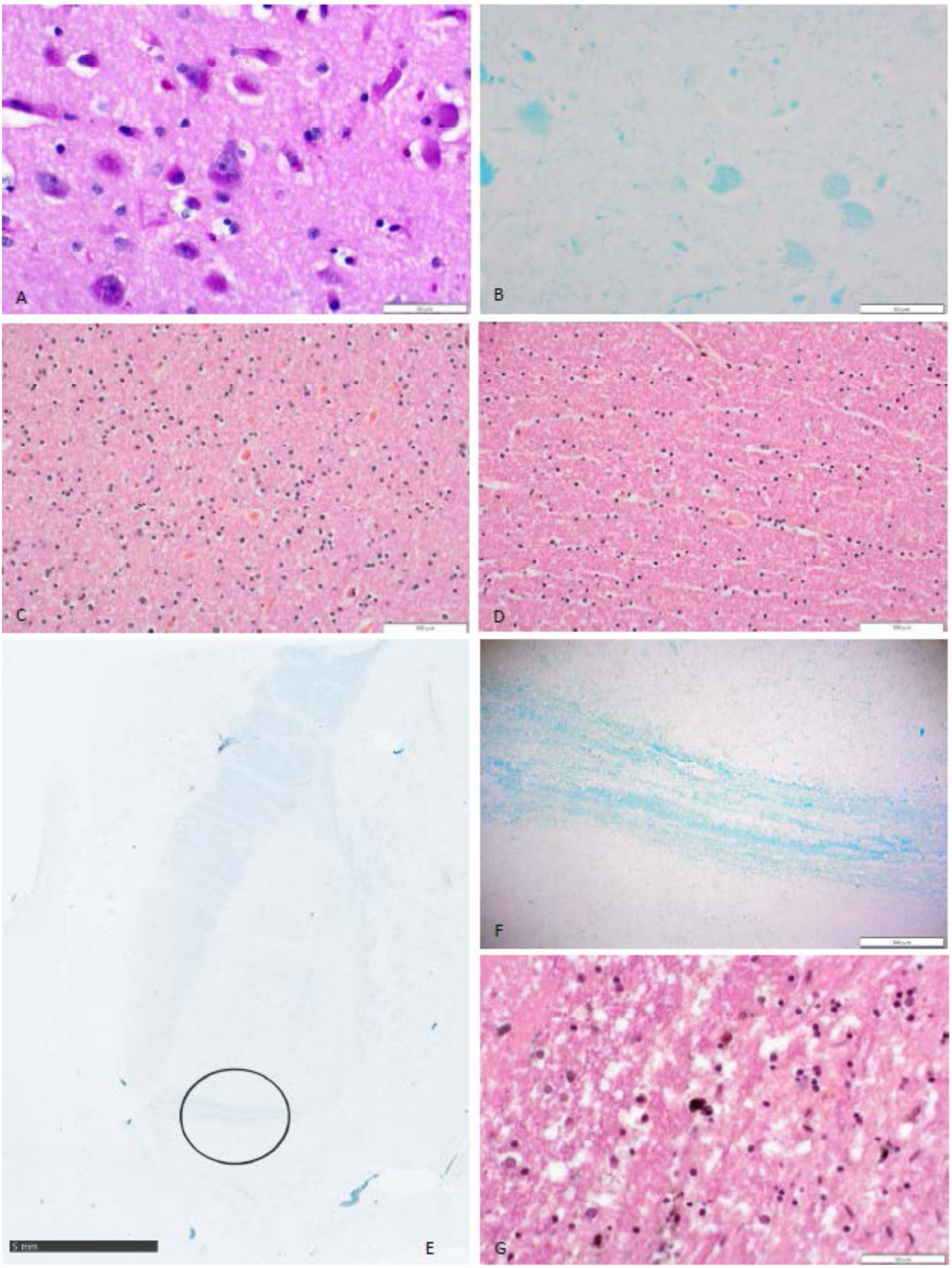
Histopathological examination of human brain tissue in MPSIIIC. **(A)** PAS-positive neuronal inclusions. (B) LFB-positive neuronal inclusions in temporal lobe neurons of patient 2. **(C)** Whiter matter hypercellularity in patient 2 compared to control **(D)**. **(E)** Focal demyelinating lesion in the anterior commissure of patient 1, seen in higher magnification in **(F)**. **(G)** Several hemosiderin-laden macrophages are adjacent to the lesion.

Further examination of the tissues of the second patient by IHC revealed that both CC and the spinal cord (SC) of the patient contained multiple CD68-positive microglia and activated GFAP-positive astrocytes consistent with pronounced neuroinflammation (Supplementary figure S5). To assess myelination, fixed tissues of CC and SC were examined by IHC using antibodies against MBP (Figure 8A and B) and MAG (Figure 9) which revealed substantially reduced levels of both markers. To determine if the levels of myelin-associated proteins are also reduced in patients affected with other subtypes of MPSIII, we analyzed PFA-fixed somatosensory cortex of post-mortem tissue, collected at autopsy and donated to the NIH NeuroBioBank. Samples of 3 MPSIII patients (MPSIIIA, MPSIIIC, and MPSIIID) and 3 non-MPS controls, matched for age and sex, were examined (project 1071, MPS Synapse). The age, cause of death, sex, race and available clinical and neuropathological information for the patients and controls are shown in Supplementary Table S1. All MPS patients had complications from their primary disease and died prematurely (before the age of 25 years). None of the patients had received enzyme replacement therapy or hematopoietic stem cell transplantation. This analysis confirmed that the amount of MBP was significantly reduced in cortices from all three MPS patients (Figure 8C), suggesting that demyelination may be a hallmark common to most subtypes of Sanfilippo disease. CC tissues from a 32-year-old MPSIIIC patient did not show any immunoreactivity for MBP or MAG, potentially due to post-mortem changes, which complicated analysis of these markers. We were also unable to achieve immunolabeling of tissues derived from a 32-year-old nor a 17-year-old patient using antibodies against CC1 and Oligo2.

**Figure 8.**
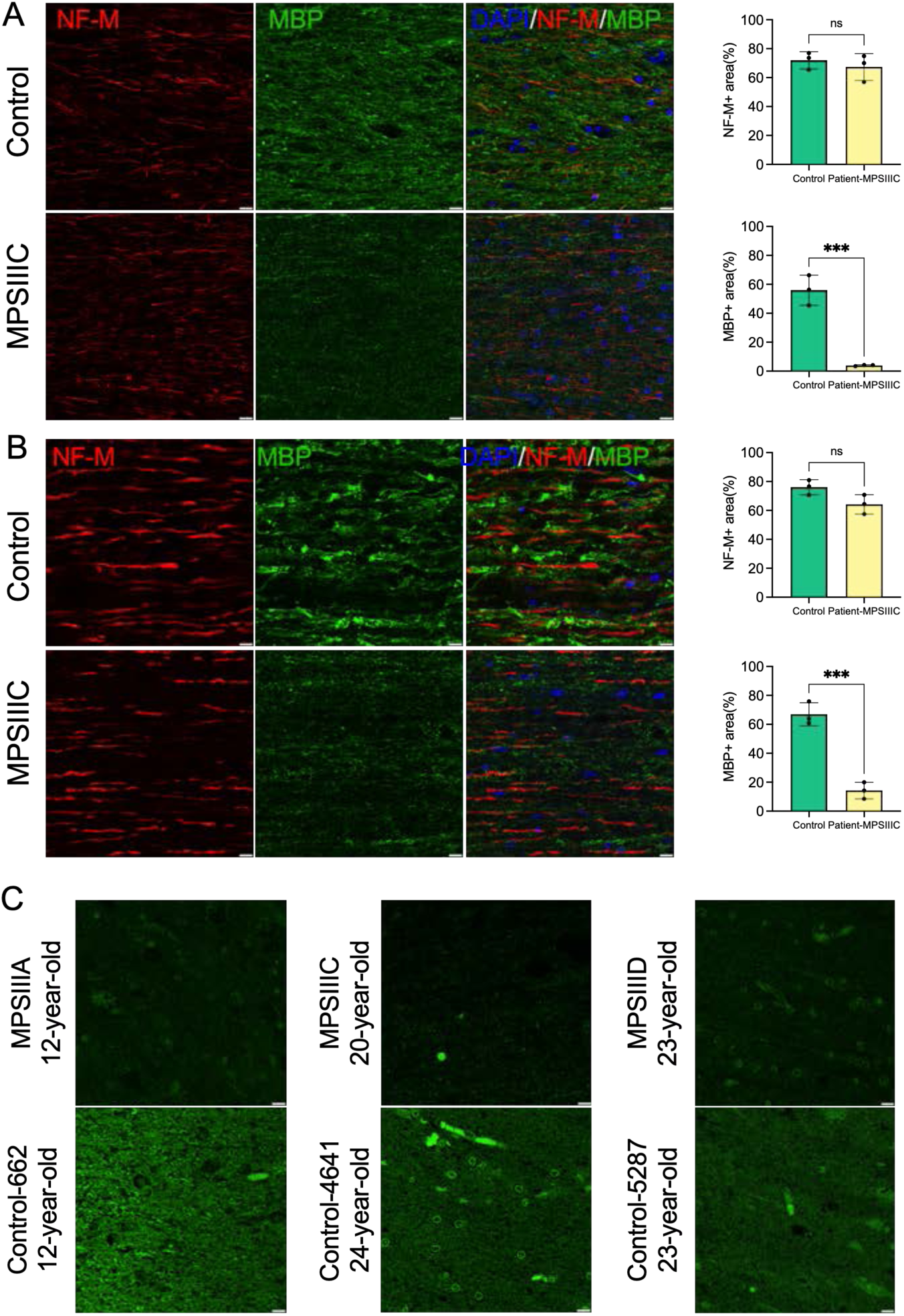
MBP levels are decreased in the brain and spinal cord of human MPSIII patients. IHC analysis reveals a significant reduction of MBP+ areas (green) on the surface of NF-M+ axons (red) in the CC **(A)** and SC **(B)** of 17-year-old MPS IIIC patient compared with age/sex matching control without a neurological disease. Scale bars equal 10 µm. Graphs show quantification of MBP+ and NF-M+ areas by ImageJ software. Individual results, means and SD of quantification performed in 3 adjacent areas are shown. P-values were calculated using an unpaired t-test. **(C)** MBP immunolabelling is also less pronounced in cortex of adult MPSIIIA, MPSIIIC and MPSIIID patients compared with age/sex matching non-MPS controls. Scale bars equal 10 µm.

**Figure 9.**
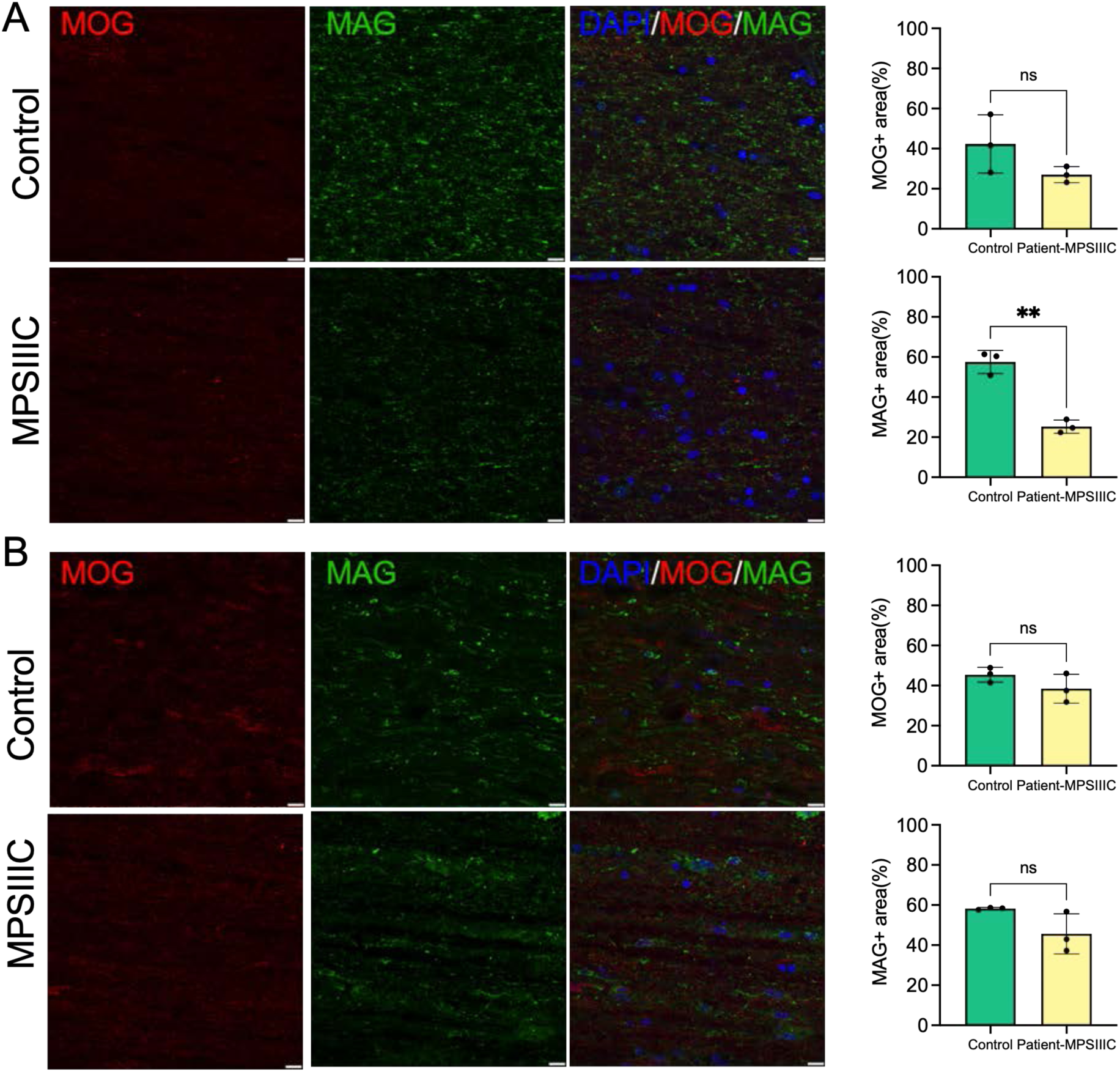
MAG levels are reduced in the CC but not in the SC of the 17-year-old MPSIIIC patient compared to control, while MOG levels show a non-significant trend toward a decrease. Panels show representative images of CC (**A**) and SC (**B**) of MPSIIIC patient and control stained with MAG+ (green) and MOG+ (red). Scale bars equal 10 µm. Graphs show quantification of MAG+ and MOG+ areas by ImageJ software. Individual results, means and SD of quantification performed in 3 adjacent areas are shown. P-values were calculated using an unpaired t-test.

Despite somewhat poor morphology of CC tissues of the 17 y/o and 32 y/o MPSIIIC patients, probably due to their post-mortem provenance, TEM analysis confirmed the presence of multiple microglia containing enlarged electrolucent vacuoles, similar to those present in the microglia in the CC of the MPSIIIC mouse brain, and consistent with storage of HS and other GAGs. OLs in the CC of the 17-year-old MPSIIIC patient also contained electrolucent vacuoles as well as the vacuoles with multilamellar inclusions, consistent with myelin accumulation (Figure 10). Multiple axons in the CC of the patients showed signs of degeneration with outfolded and split myelin containing cytoplasmic materials inside the sheaths. Axonal swellings containing vesicles, similar to those present in the MPSIIIC mice, were also detected. These structural abnormalities were not observed in the CC of the age and sex-matched non-MPS patient.

**Figure 10.**
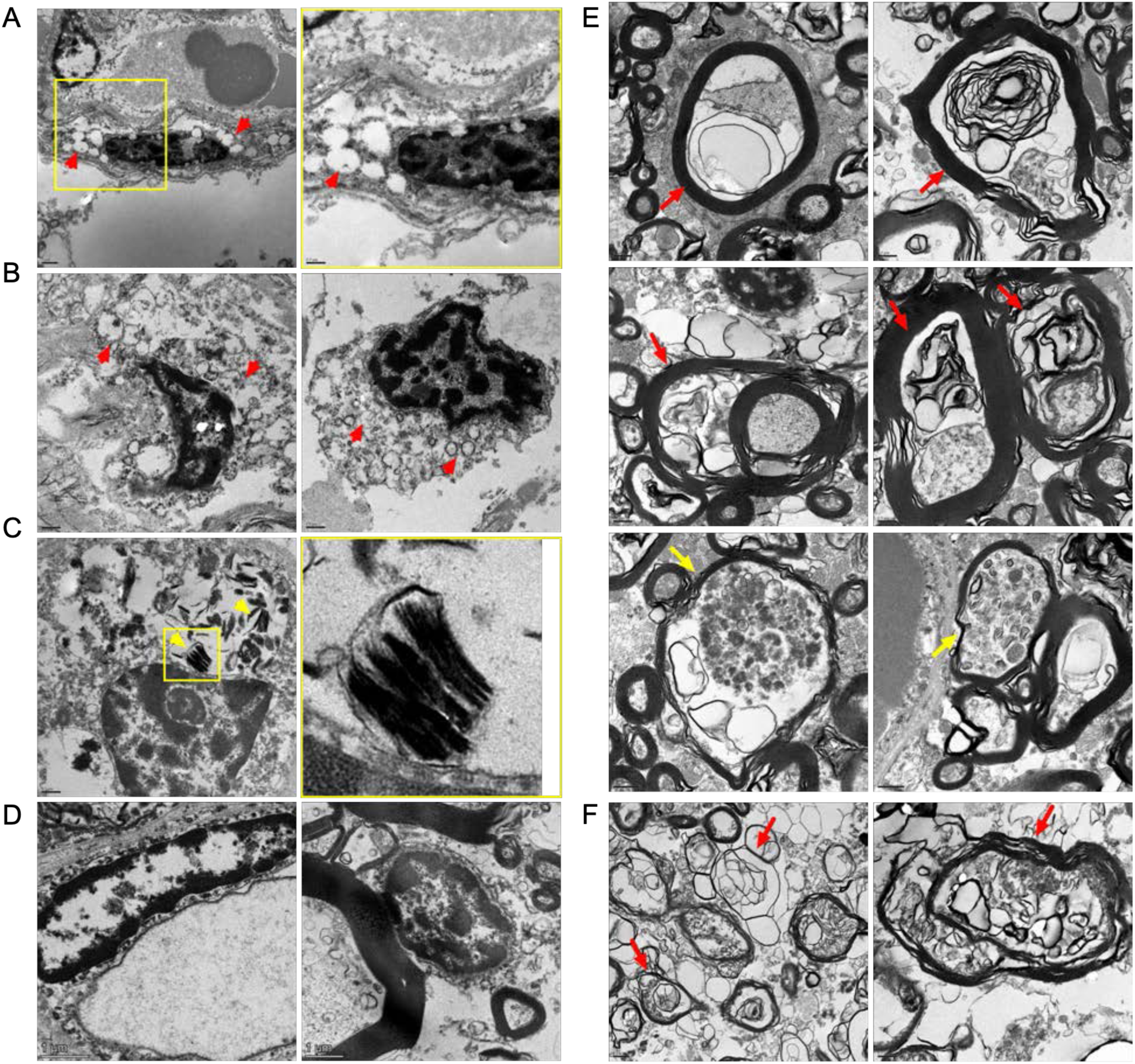
TEM analysis confirms microgliosis, pathological changes in oligodendrocyte morphology and axonopathy in the CC of MPSIIIC patients. Electron micrographs show electron-lucent vacuoles (red arrowheads) consistent with HS storage in microglia of the CC of the 17-year-old (**A**) and 32-year-old (**B**) MPSIIIC patients. Oligodendrocytes in the CC of the 17-year-old MPSIIIC patient contain zebra bodies consistent with myelin accumulation (**C**, yellow arrowheads). Microglia (left panel) and oligodendrocytes (right panel) in the CC of control do not contain storage vacuoles (**D**). Degenerated axons with outfolded and split myelin, sometimes containing cytoplasmic pockets between the sheaths (red arrows), as well as large axonal swellings containing accumulating vesicles (yellow arrows), are observed in the CC of the 17-year-old (**E**) and 32-year-old (**F**) MPS IIIC human patients. These structural abnormalities are not observed in the CC of the age and sex-matched non-MPS patient.

## Discussion

Here we identify persistent demyelination as a key component of CNS pathology in MPSIIIC, and likely associated with all diseases of the MPSIII spectrum. First, we determined that the amount of MBP, a protein marker of myelinated axons, was drastically reduced in CC of both MPSIIIC mice and MPSIIIC patients at advanced stages of the disease, consistent with myelin disruption and demyelination. Other myelin markers, MAG and MOG were also reduced. Second, ultrastructural analysis of brain tissue confirmed that the myelin sheaths in the MPSIIIC brains were reduced in thickness and revealed structural abnormalities, including outfolded, empty or split myelin. Third, regions of white matter in MPSIIIC mouse brain were massively infiltrated by activated CD68-positive microglia with lysosomal accumulation of MBP-positive elements and multilamellar fragments (“zebra bodies”), consistent with phagocytosis of myelin fragments. Occurrence of similar processes associated with demyelination and axonal degeneration have been reported in multiple white matter pathologies including ischemic injuries, multiple sclerosis and other inflammatory demyelinating diseases, optic nerve diseases, aging, and experimental models of immune encephalomyelitis (reviewed in [44–46]). In immune encephalomyelitis, activated proinflammatory microglia were reported to directly trigger the loss of myelin, however in MPSIIIC mice, high levels of microgliosis is observed starting from a very early age (P25 or even before), while the loss of MBP was not detected before the age of 6 months. These findings support the conclusion that axonal demyelination in MPSIIIC is due to aberrant myelin maintenance by resident OLs rather than defects of early axonal development or damage caused by activated microglia or astrocytes. This was supported by our observation that OLs in the CC of MPSIIIC mice were scarce and, in general, immature. Further analysis of OL morphology revealed that they contained multiple electrolucent vacuoles, consistent with storage of HS, and multilamellar bodies, characteristic of lipid accumulation. This was confirmed by IHC, demonstrating that OLs in MPSIIIC but not WT mice were positive for HS and GM3 ganglioside. Based on the presence of these morphological abnormalities, we speculate that the majority OLs in MPSIIIC mice are either dysfunctional or have reduced functionality.

We and others have previously demonstrated that the levels of GM3 ganglioside, together with GM2 ganglioside, lactosylceramide and glucosylceramide, are drastically increased in the brains of mouse models of MPSIII and in the brains of MPSIII human patients [35, 36, 47]. The accumulation of these secondary materials mainly occurs in pyramidal neurons in the deep cortex layers and in the CA1-CA3 regions of the hippocampus [48, 49]. Our current data show that in contrast to other brain cells, OLs in the CC solely accumulate GM3, while rare GM2-positive cells scattered across the CC are negative for the OL marker Olig2. Although the mechanism underlying the massive accumulation of GM3 ganglioside in the OLs of MPSIIIC brain requires further investigation, we are tempted to speculate that this phenomenon may be related to the development of myelin defects. Previous studies demonstrated that in Niemann-Pick type C (NP-C) the accumulation of GM3 in OLs is a prerequisite for dysfunction and demyelination [13]. Similarly to MPSIII, NP-C manifests with predominant storage of GM2 and GM3 gangliosides [50] and myelin defects in the brain tissues (Reviewed in [3]). In the mouse model of NP-C (*Npc1^−/−^* mice) hypomyelination is pronounced in the cerebral cortex and the CC, presumably accounting for the tremor and ataxia observed in these animals [51]. The *Npc1^−/−^* mice, heterozygous for the deletion of the *Siat9* (GM3 synthase) gene, showed ameliorated neuropathology, including motor disability and demyelination [13]. At the same time, deletion of *Siat9* gene resulted in the enhanced neuropathological phenotype and demyelination in *Npc1^−/−^*mice, indicating that the presence (but not an excessive storage) of GM3 ganglioside is essential for OLs development [13].

Importantly, OLs in the CC of MPSIIIC mice also exhibited mitochondrial pathology; the majority of mitochondria in these cells had a dysmorphic ballooned appearance with absent cristae. Similar pathology has been previously described by our team in the hippocampal pyramidal neurons of MPSIIIC mice, which was associated with a progressive loss of mitochondrial activity and neurodegeneration [38]. In the adult brain, once myelination is complete, the long-term integrity of axons is known to depend on the supply of energy to myelinating cells that are key for preserving the connectivity and function of a healthy nervous system (Reviewed in [52]). Thus, it is tempting to speculate that mitochondrial dysfunction in OLs, associated perhaps with reduced maturation and increased degeneration, may underlie the demyelination that occurs in the MPSIIIC brain.

Major findings in the mouse MPSIII model were confirmed with the brain tissues of human MPSIIIC patients, suggesting that demyelination and white matter injury contribute to the pathology at least at the later stages of the disease. Importantly, LFB staining did not reveal regions with profound demyelination in the brains of both patients, except for one region in the brain of a 38-year-old patient, where the loss of LFB staining was associated with the presence of hemosiderin-laden macrophages suggesting an occurrence of an old micro hemorrhage. This may explain, why myelination defects were not previously described in the majority of pathology reports where brain tissues of MPSIII patients have been examined using only traditional histochemical techniques such as HES, PAS and LFB staining. Further studies are required to determine whether patients affected with all subtypes of Sanfilippo disease show a similar degree of demyelination. We consider this to be plausible, as drastically reduced levels of MBP staining were detected in the post-mortem cortical samples of MPSIIIA and MPSIIID patients obtained from NeuroBiobank.

Magnetic resonance diffusion tensor imaging allows generation of quantitative maps that describe tissue microstructure by decomposing diffusion signals into axial diffusivity, radial diffusivity, and fractional anisotropy. We have previously shown how a reduction in radial diffusivity can reliably quantify myelination during development [53]. Consistent with this, increased radial diffusivity has been used as a biomarker of demyelination [40, 41, 54]. A more recent method, Diffusion Basis Spectrum Imaging can provide additional information for an extra-axonal compartment, such as fiber fraction, hindered diffusivity (extracellular), restricted diffusivity (intracellular) and water fraction [30]. Restricted diffusivity has, in particular, been shown to reflect inflammation with increased cellularity in a mouse model of neuroinflammation [55]. It has been also applied to detect white matter injury with edema or tissue loss [56–58]. Interestingly, our analysis of diffusivity maps of the CC of 7-month-old MSCIIIC mice revealed a significant increase in radial diffusivity, suggesting loss of myelin, together with a strong increase of mid-higher isotropic diffusivity indicative of edema and tissue loss. In line with the absence of axonal loss, our MRI analysis found no changes in fiber fraction, fractional anisotropy or axial diffusivity. On the other hand, we did not detect significant changes in restricted diffusivity, which was expected considering the high level of microgliosis in the CC of MPSIIIC mice. It is possible, however that, the model we used was not sufficiently sensitive to detect subtle increase in local cellularity because of the very dense structure of white matter mainly composed of tightly organized fibers.

Together, our data provide novel insights into pathophysiology of Sanfilippo disease. They also identify widespread demyelination in CNS as an important biomarker of disease progression and suggest that analysis of brain myelin by MRI may become in the future a leading non-invasive method for clinical patient assessment.

## Abbreviations

APC(CC-1): Adenomatous polyposis coli (APC) clone CC1
ARSA: Arylsulfatase A
BSA: Bovine serum albumin
CC: Corpus callosum
CCAC: Canadian Council on Animal Care
CNS: Central Nervous System
CT-scan: Computed tomography-scan
DBSI: Diffusion Basis Spectrum Imaging
DTI: Diffuse tension imaging
EGFP: Enhanced green fluorescent protein
EM: Electron microscopy
GAG: Glycosaminoglycan
GALC: β-galactocerebrosidase
GNS: N-acetylglucosamine-6-sulfate sulfatase
HGSNAT: Heparan sulphate acetyl-CoA: α-glucosaminide N-acetyltransferase
HS: Heparan sulphate
IHC: Immunohistochemistry
KI: Knock-in
LAMP: Lysosomal associated membrane protein
LSD: Lysosomal storage disorder
MAG: Myelin-associated glycoprotein
MBP: Myelin basic protein
MLD: Metachromatic leukodystrophy
MOG: Myelin oligodendrocyte glycoprotein
MPS: Mucopolysaccharidosis
MRI: Magnetic resonance imaging
NAGLU: N-acetyl-α-D-glucosaminidase
NF-M: Neurofilament medium chain
NPC: Niemann-Pick disease type C
OCT: Optimum cutting temperature
OL: Oligodendrocyte
Oligo2: Oligodendrocyte transcription factor2
PFA: Paraformaldehyde
SC: Spinal cord
SGSH: N-sulfoglucosamine sulfohydrolase
SSC: Somatosensory cortices
TEM: Transmission electron microscopy
WT: Wild type

