## Supplementary for "Severe Central Nervous System Demyelination in Sanfilippo Disease"

**Supplementary data**

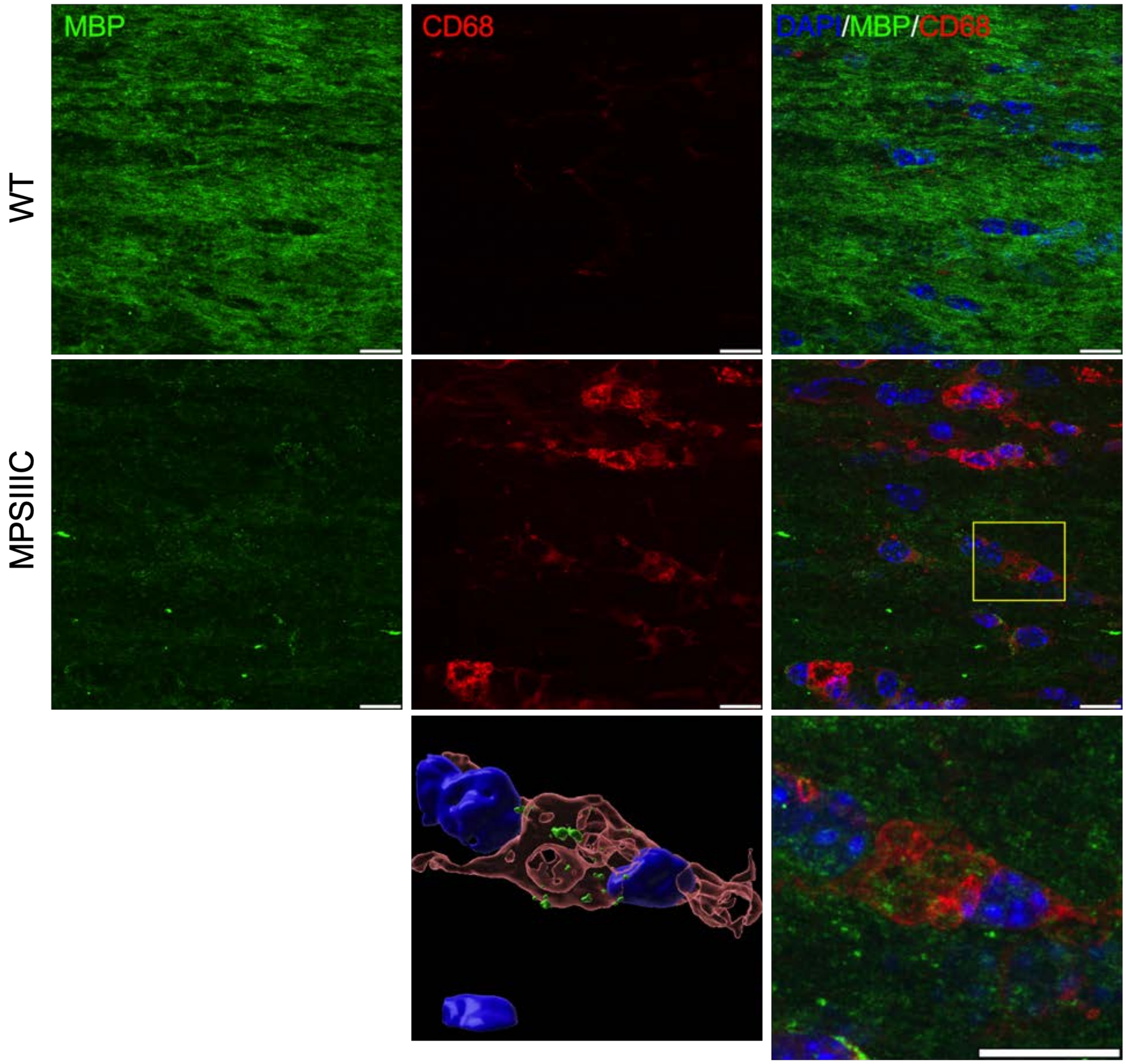


**Figure S1. Intracellular aggregation of MBP in microglia in CC of MPSIIIC mice.**

Panels show representative confocal microscopy images of CC tissue of 6-month-old WT and MPSIIIC mice labelled with antibodies against MBP (green) and CD68 (red). DAPI (blue) was used as a nuclear counterstain. Scale bars equal 10 µm. The enlarged image of the boxed area shows a CD68+ activated microglia containing MBP+ puncta. 3D reconstruction shows that MBP+ puncta are located inside the microglia cell.

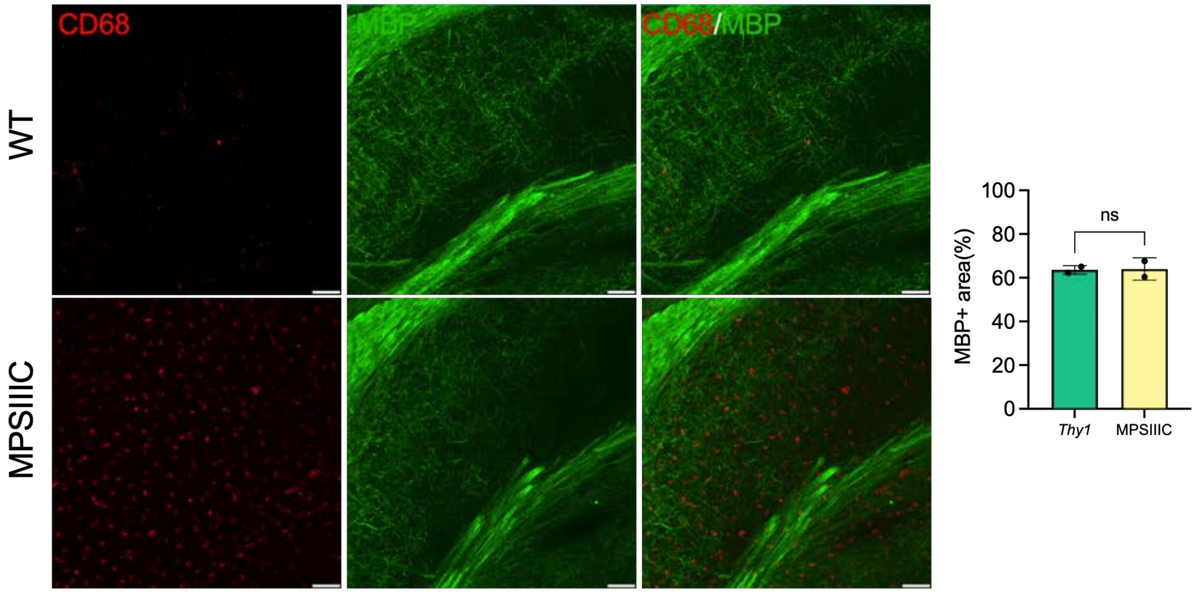


**Figure S2. Microgliosis does not coincide with the loss of myelin at an early age.**

Panels show representative confocal microscopy images of hippocampal and CC tissues of P25 WT and MPSIIIC mice labelled with antibodies against CD68 (red) and MBP (green). Scale bars equal 50 µm.

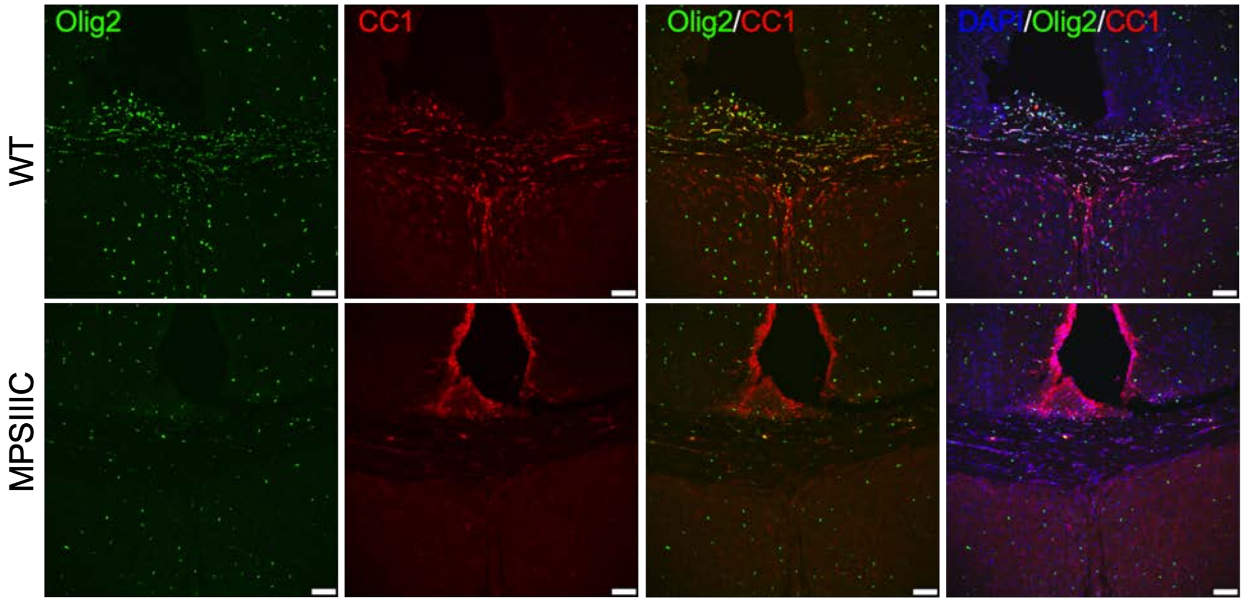


**Figure S3. CC of MPSIIIC mice show reduced numbers of mature oligodendrocytes.**

Panels show representative images of the CC of 6-month-old WT and MPSIIIC mice immunolabelled for OL lineage marker Olig2 (green) and mature OL marker CC1 (red). DAPI (blue) was used as a nuclear counterstain. Scale bar equals 50 mm.

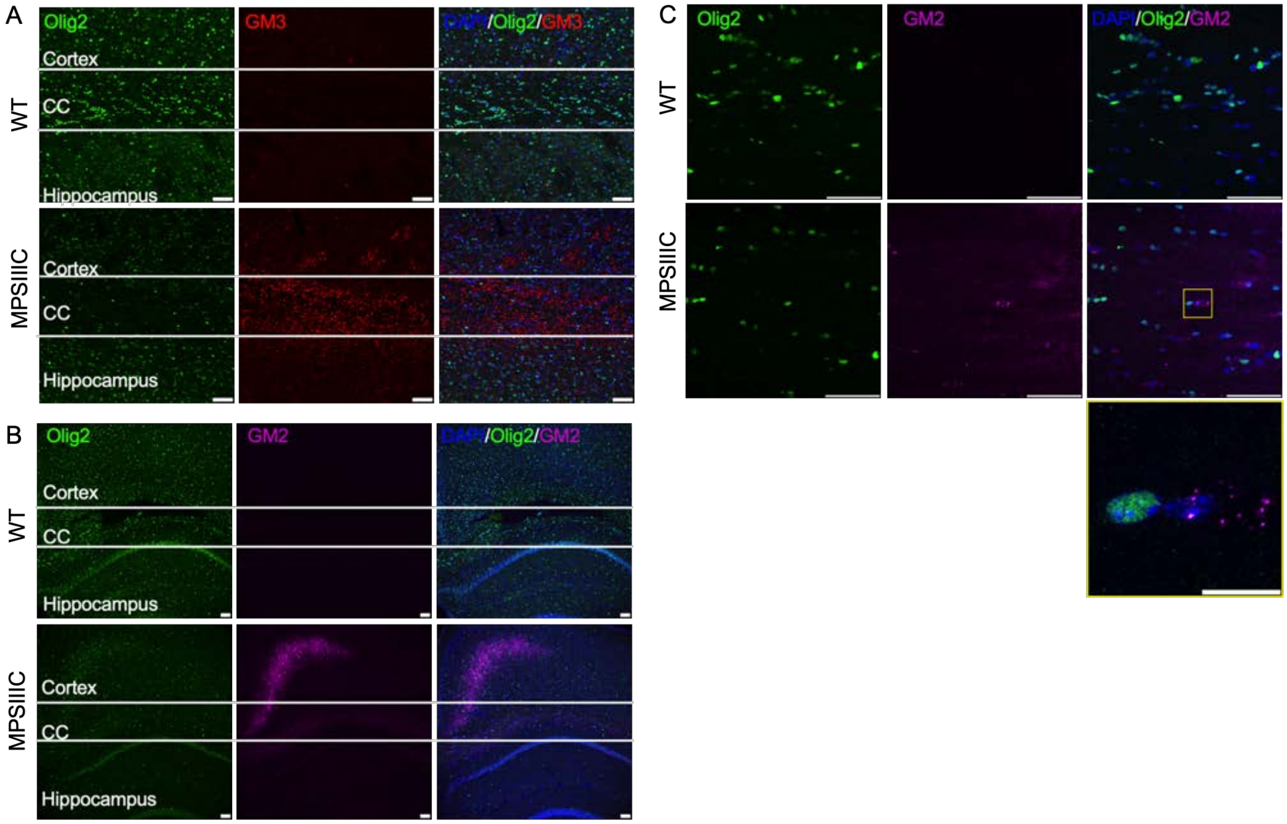


**Figure S4. Secondary storage of GM3 and GM2 gangliosides in the brain of MPSIIIC mice**.

Panels show confocal microscopy images of the cortex, CC and hippocampus tissues of 6-month-old WT and MPSIIIC mice stained with antibodies against Olig2 (green), GM3 ganglioside (red) **(A)** and GM2 ganglioside (purple) **(B),** revealing storage of GM3 ganglioside in the CC and GM2 ganglioside in the cortex of MPSIIIC but not of WT mice. **(C)** GM2 ganglioside does not accumulate in OLs in CC tissue of 6-month-old WT and MPSIIIC mice. DAPI (blue) was used as a nuclear counterstain. Scale bar equals 50 mm for **A, B, C,** and 10 mM for the zoomed image in the panel **C**.

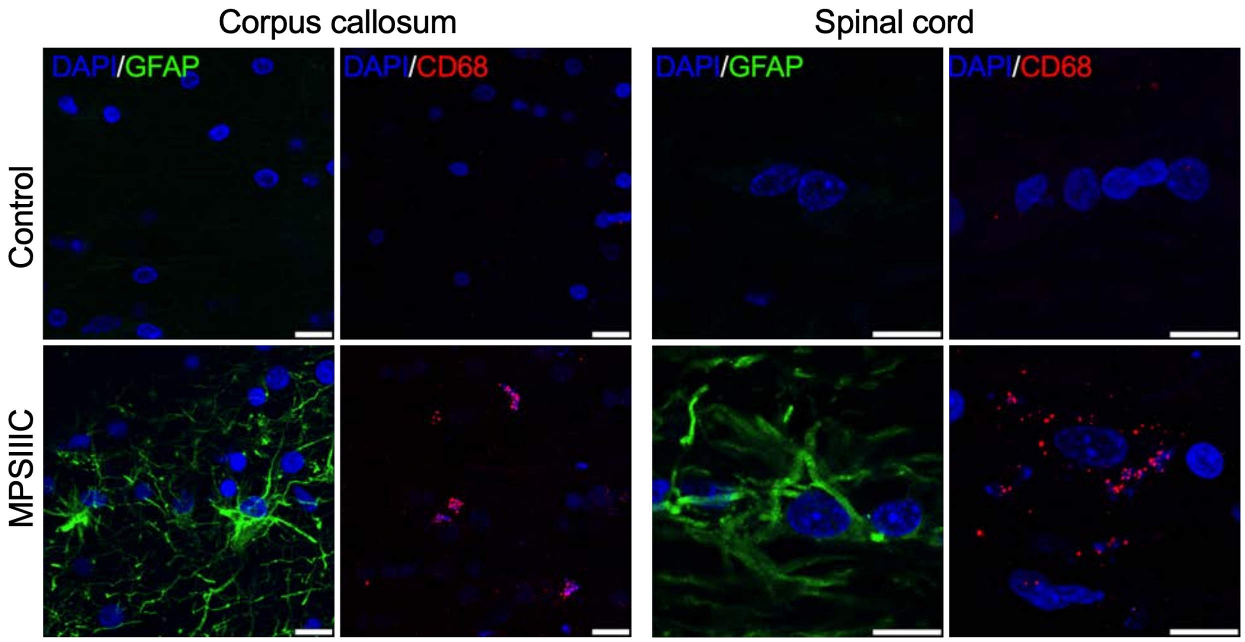


**Figure S5. Pronounced astromicrogliosis in the corpus callosum and spinal cord of a 17-years-old MPSIIIC patient.** Multiple GFAP-positive astrocytes (green) and C68-positive microglia (red), indicative of neuroimmune response, are detected in the CC and SC of the MPSIIIC patient but not of the age/sex matching non-MPS control.
